# Modulating inter-mitochondrial contacts to increase membrane potential for mitigating blue light damage

**DOI:** 10.1101/2025.10.24.684455

**Authors:** Yuxin Wang, Kangqiang Qiu, Weiwei Zou, Prativa Amom, Tushar H. Ganjawala, Eugene Lee, Zhiqi Tian, Xiuqiong Xu, Taosheng Huang, Nien-Pei Tsai, Donglu Shi, Ping Kang, Hua Bai, Amanda L. Zacharias, Kai Zhang, Jiajie Diao

**Affiliations:** Department of Cancer Biology, University of Cincinnati College of Medicine, Cincinnati, OH 45267, USA; The Materials Science and Engineering Program, Department of Mechanical and Materials Engineering, University of Cincinnati, Cincinnati, OH 45221, USA; Division of Human Genetics, Cincinnati Children’s Hospital Medical Center, Cincinnati, OH 45229, USA; Divisions of Developmental Biology and Neonatology and Pulmonary Biology, Cincinnati Children’s Hospital Medical Center, Cincinnati, OH 45229, USA; Department of Pediatrics, University at Buffalo, 1001 Main Street, Buffalo, NY 14203, USA; Department of Molecular & Integrative Physiology, School of Molecular and Cellular Biology, University of Illinois at Urbana-Champaign, Urbana, IL 61801, USA; Department of Biomedical Engineering College of Engineering and Applied Science, University of Cincinnati, Cincinnati, OH 45221, USA; Department of Genetics, Development, and Cell Biology, Iowa State University, Ames, IA 50011, USA; Department of Pediatrics, University of Cincinnati College of Medicine, Cincinnati, OH 45267, USA; Department of Biochemistry, School of Molecular and Cellular Biology, University of Illinois at Urbana-Champaign, Urbana, IL 61801, USA; Science and Technology Center for Quantitative Cell Biology, University of Illinois at Urbana-Champaign, Urbana, IL 61801, USA

**Keywords:** membrane contact sites, mitochondrial membrane potential, mitochondria, optogenetics

## Abstract

Mitochondrial membrane potential (MMP) is essential for mitochondrial functions, yet current methods for modulating MMP lack precise spatial and temporal control. Here, we developed an optogenetic mitochondrial tethering system, termed OptoMitoTether, that enables reversible control of inter-mitochondrial proximity with high spatiotemporal precision. Blue light activation induced CRY2PHR oligomerization between adjacent mitochondrial outer membranes, reduced mitochondrial nearest-neighbor distances, and promoted inter-mitochondrial tethering. OptoMitoTether activation was associated with elevated MMP and was associated with lower reactive oxygen species (ROS) and preserved ATP production during prolonged blue light stress. Moreover, OptoMitoTether expression preserved mitochondrial bioenergetic function in human retinal cells and extended lifespan in *C. elegans* during chronic blue light exposure. These findings establish OptoMitoTether as a tool for manipulating mitochondrial proximity and reveal a link between inter-mitochondrial tethering and mitochondrial bioenergetic state during blue light stress.

## Introduction

Mitochondria, often referred to as the ‘powerhouse of the cell’, are essential organelles responsible for ATP production and play a critical role in regulating key cellular processes (Alshaabi et al., 2020; Bulthuis et al., 2019), including calcium signaling, nutrient sensing, innate immune responses, and programmed cell death (Chen et al., 2023; Lim et al., 2021; Martínez-Reyes and Chandel, 2020; Mills et al., 2017; Rambold et al., 2011). The mitochondrial membrane potential (MMP) provides the electrochemical proton gradient that drives ATP synthesis, serving as an intermediate form of stored energy during oxidative phosphorylation (L D Zorova et al., 2018; Ljubava D Zorova et al., 2018). Beyond its energetic role, MMP is also essential for maintaining mitochondrial and cellular homeostasis through non-energetic functions, including ion transport, protein import, and mitophagy (Cunha et al., 2011; Korshunov et al., 1997; Miller and Sheetz, 2004; Novak, 2012; Paschen and Neupert, 2001; Sato et al., 2019; Scott et al., 1980; Youle and Narendra, 2010).

However, stimulus and stressors, such as excessive reactive oxygen species, can disrupt mitochondrial dynamics by promoting fragmentation, disturbing mitochondrial contacts, impairing electron transport, and causing MMP depolarization and heterogeneity (Kuznetsov and Margreiter, 2009; Picard et al., 2011). Depolarization refers to a reduction in the inner mitochondrial membrane potential, leading to a weakened proton motive force and diminished ATP synthesis capacity. In contrast, heterogeneity refers to the variability in MMP among mitochondria, reflecting spatial bioenergetic imbalance and functional asymmetry within the mitochondrial network. The disruption or loss of MMP has been linked to pathological conditions, including neurodegenerative disorders, metabolic syndromes, and cardiovascular diseases, where impaired mitochondrial communication drives cellular dysfunction (Chen et al., 2007; Dorn, 2015; Pokharel et al., 2024; Shah et al., 2019; Suárez-Rivero et al., 2016; Szendroedi et al., 2011; Wang et al., 2009; Yapa et al., 2021). Ischemia-reperfusion events frequently trigger prolonged opening of the mitochondrial permeability transition pores, leading to abrupt loss of MMP and widespread mitochondrial dysfunction (Halestrap and Richardson, 2015; Hausenloy and Yellon, 2004). Similarly, aging-related accumulation of mitochondrial DNA damage and a decline in mitophagy efficiency lead to a progressive decrease in MMP and increased heterogeneity, ultimately compromising mitochondrial bioenergetics in aged tissues. Persistent MMP depolarization impairs ATP production, disrupts calcium homeostasis, and increases susceptibility to apoptosis due to compromised mitochondrial integrity and signaling (Green and Reed, 1999; Hüttemann et al., 2008; Ly et al., 2003; Santo-Domingo and Demaurex, 2010). Moreover, elevated MMP heterogeneity has been linked to inefficient oxidative phosphorylation and metabolic imbalance, contributing to tissue dysfunction and disease progression (Glancy and Balaban, 2012; Rossignol et al., 2004; Rovini et al., 2021).

Because MMP is essential for mitochondrial functions, restoring MMP would be critical to maintain energy supply for cells. Indeed, several classes of interventions have been developed to preserve or restore MMP under stress conditions. One widely used approach targets the mitochondrial permeability transition pore (mPTP). For example, ischemia-reperfusion injury causes a sudden collapse of MMP due to prolonged opening of mPTP (Halestrap and Richardson, 2015). Uncoupling inhibitors like cyclosporin A inhibit mPTP opening, thereby preventing catastrophic proton leakage and membrane depolarization. However, their effects are context-dependent and limited to conditions where mPTP activation is a primary driver of MMP loss. A second strategy involves mitochondria-targeted antioxidants, such as MitoQ, which aim to neutralize excessive reactive oxygen species (ROS) within mitochondria (Smith et al., 2011). ROS accumulation is a major contributor to electron transport chain (ETC) dysfunction and MMP instability. Nonetheless, efficacy of antioxidants can be compromised by uneven ROS distribution or irreversible oxidative damage, and systemic delivery often leads to off-target effects. A third approach employs metabolic regulators, including nicotinamide riboside (NR) and coenzyme Q10, which boost cellular NAD levels or improve ETC substrate availability (Gomes et al., 2013; Trammell and Brenner, 2013). These agents enhance mitochondrial respiration and indirectly stabilize MMP. Yet, they do not act on the membrane potential directly and are limited by variable cellular uptake, short half-lives, and the inability to restore MMP when structural mitochondrial damage is present. Collectively, these pharmacological interventions offer partial and often transient rescue for mitochondrial function. These approaches primarily target biochemical pathways and do not provide direct spatiotemporal control over mitochondrial organization or inter-mitochondrial proximity.

Meanwhile, the dynamic spatial organization of mitochondria also mediates mitochondrial functions by forming interconnected networks that enable mitochondria to exchange metabolites, buffer calcium fluctuations, and dynamically adapt to cellular demands. Beyond fusion and fission, the spatial organization of neighboring mitochondria and their physical apposition may provide an additional layer of mitochondrial coordination (Chen and Chan, 2009; Detmer and Chan, 2007; Duchen, 2000; McBride et al., 2006; Nunnari and Suomalainen, 2012; Szabadkai et al., 2004). Emerging evidence suggests that physical proximity between distinct membranes could influence membrane potential (Bu et al., 2016; Gurunian et al., 2025). Inter-mitochondrial junctions have been associated with coordinated cristae organization and functional coupling between neighboring mitochondria. In addition, nanoscale apposition of charged membranes or proteins can alter the local electrostatic environment and membrane potential (Picard et al., 2015). Together, these observations raise the possibility that controlling mitochondrial proximity, instead of directly manipulating the canonical fusion and fission machinery, could provide a means to modulate mitochondrial bioenergetics.

Furthermore, mitochondria experience a range of cellular stresses. With the widespread integration of blue light, a predominant component of light-emitting diodes (LEDs), into modern life, human exposure to high-energy light has significantly increased, raising concerns about new forms of cellular damage. Indeed, excessive exposure to blue light can cause mitochondrial damage, contributing to serious ocular disorders (Li et al., 2018; Tao et al., 2019; Wang et al., 2023; Yan et al., 2025). At the subcellular level, blue light exposure has been shown to disrupt mitochondrial morphology, compromise functional integrity, and reduce MMP (Godley et al., 2005; Hao et al., 2023; Tao et al., 2019). Blue light therefore provides a unique context for testing a stimulus-coupled optogenetic protection strategy, in which the same environmental input that causes damage during prolonged exposure also activates the protective mitochondrial tethering system.

In this study, we developed an optogenetic mitochondrial tethering system, OptoMitoTether, to physically reduce the separation between distinct mitochondria and induce reversible tethering. We found that this optogenetic modulation alters MMP, ROS production, and bioenergetic function under short-term activation and prolonged blue light stress. Furthermore, the activation of OptoMitoTether exerts protective effects in human retinal cells and *C. elegans*. As the OptoMitoTether is activated exclusively by blue light, it provides an *in situ* strategy to mitigate blue light-induced mitochondrial damage. This approach offers valuable potential for both fundamental studies and therapeutic interventions in ophthalmic and other mitochondrial dysfunction-related diseases.

## Results

### OptoMitoTether induces reversible inter-mitochondrial tethering

Inspired by previous studies indicating that nanoscale proximity of a charged membrane or protein condensate to a membrane amplifies the local membrane potential (Bu et al., 2016; Gurunian et al., 2025), we developed CRY2PHR-mCherry-Miro1TM, hereafter termed OptoMitoTether, to bring mitochondrial membranes into contact by light in living cells (Figure 1a). The photolyase homology region of cryptochrome (CRY2PHR) (Bugaj et al., 2015; Chang et al., 2014; Che et al., 2015; Park et al., 2017; Taslimi et al., 2014; Zhang and Cui, 2015) was anchored to the outer mitochondrial membrane via the mitochondria-targeting transmembrane domain of Miro1 (Miro1TM) (Duan et al., 2015). Upon blue light stimulation (300 μW/cm^2^), CRY2PHR undergoes oligomerization, which brings their bound mitochondria to proximity to form inter-mitochondrial tethering. During this process, the sites of CRY2PHR oligomerization exhibit higher fluorescent signals at light-induced contact sites. Structured illumination microscopy (SIM) (Figure S1) revealed that the red fluorescence initially delineated mitochondrial morphology but subsequently formed puncta upon light-activation in less than 5 min (Figure S2).

**Figure 1.**
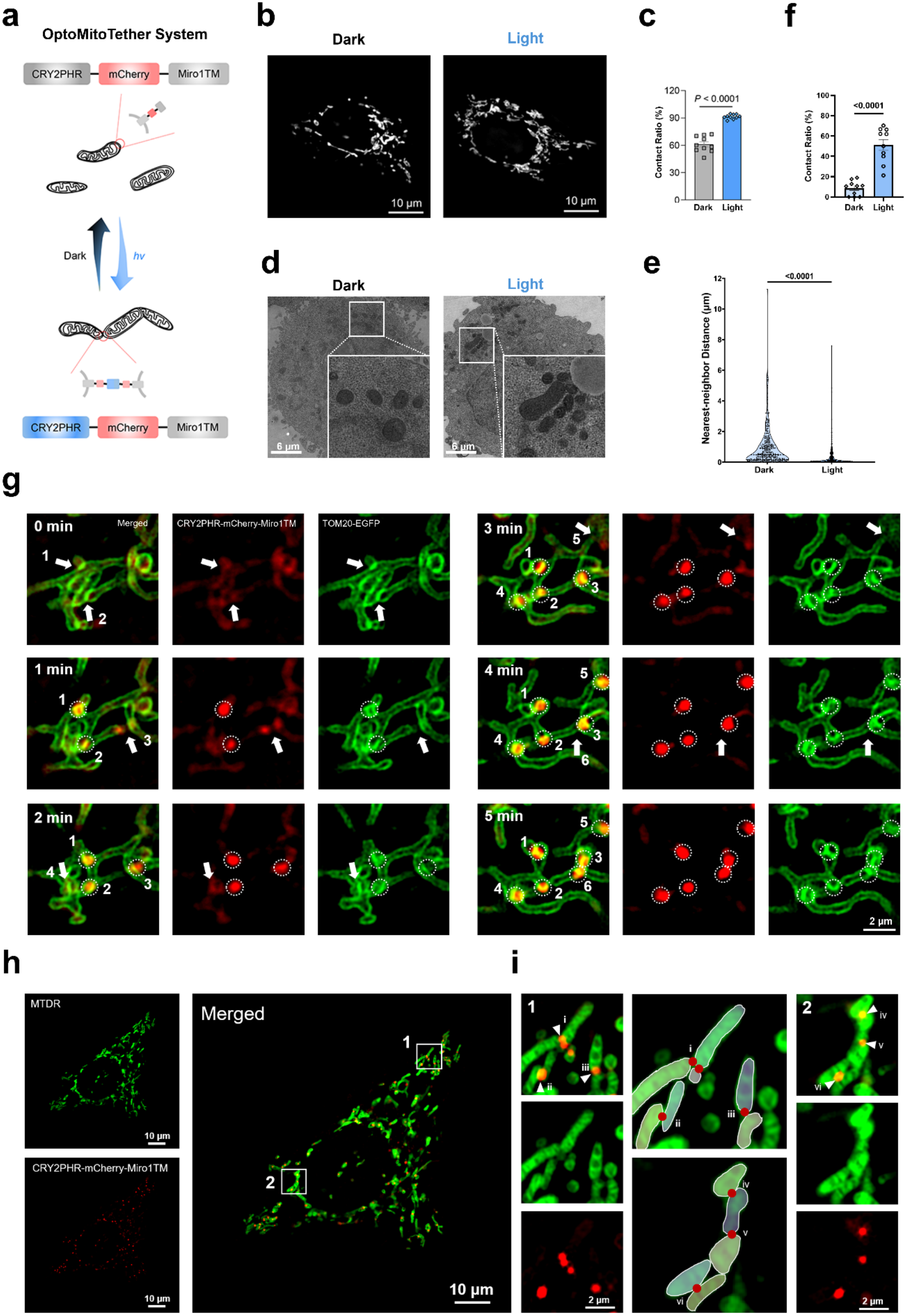
OptoMitoTether induces reversible inter-mitochondrial tethering. **(a)** Schematic representation of the blue light induced mitochondrial contacts. The light-sensitive proteins CRY2PHR were anchored to mitochondria via the specific organelle-targeting transmembrane domain Miro1TM. Under blue light illumination, the CRY2PHR interacted with each other to form mito-contact. Meanwhile, the fluorescent protein mCherry served as a marker of the plasmid’s expression. **(b)** Representative SIM images of OptoMitoTether-expressing HeLa cells under dark conditions and following 20 min of blue light illumination (300 μW/cm²). Mitochondria were visualized with MTG. **(c)** The percentage contact ratio of mitochondria from SIM images of HeLa cells corresponding to (b). Data are given as mean ± SEM (n = 10). P < 0.0001, two-tailed Student’s t test. **(d)** Representative TEM images of OptoMitoTether-expressing HeLa cells under dark conditions and following blue light activation. Insets show enlarged examples of neighboring mitochondria. **(e)** Distribution of nearest-neighbor mitochondrial distances measured from TEM images under dark and light conditions. For each mitochondrion, the shortest outer-membrane-to-outer-membrane distance to the nearest distinct mitochondrion was measured. P < 0.0001, nested t test. **(f)** TEM-defined mitochondrial contact ratio under dark and light conditions. A TEM contact was defined as an outer-membrane-to-outer-membrane distance <25 nm. Each point represents one cell. Data are mean ± SEM; n = 10 cells. P < 0.0001, two-tailed Student’s t test. **(g)** Time-lapse SIM imaging of a HeLa cell co-expressing TOM20-EGFP and CRY2PHR-mCherry-Miro1TM during 0–5 min of blue light activation. TOM20-EGFP delineates mitochondrial outer-membrane profiles (green), whereas CRY2PHR-mCherry is shown in red. Numbered circles indicate the successfully formed CRY2 aggregates (puncta) during illumination, and arrows mark the corresponding pre-existing interfaces where new aggregates are about to emerge in the subsequent frame. Scale bar, 2 μm. **(h)** Representative pseudocolored SIM image showing mitochondrial morphology and CRY2PHR-mCherry enrichment following OptoMitoTether activation. MTDR is displayed in green and CRY2PHR-mCherry in red for visualization. **(i)** Enlarged examples of representative tip-to-tip, side-by-side, and tip-to-side mitochondrial appositions from (h). CRY2PHR-mCherry-enriched puncta are shown in red, and schematic overlays illustrate the geometry of the corresponding mitochondrial profiles.

To better visualize the change of mitochondria before and after optogenetic stimulation, living HeLa cells expressing CRY2PHR-mCherry-Miro1TM were subsequently stained with either MitoTracker^TM^ Deep Red (MTDR) or MitoTracker^TM^ Green (MTG) before and after 20 minutes of blue light exposure. As illustrated in Figure 1b, blue light irradiation significantly increases the interaction between mitochondria. Blue light activation increased the proportion of mitochondria exhibiting close apposition with at least one neighboring mitochondrion from 60.79% to 91.34% (Figure 1c). We next quantified mitochondrial proximity using transmission electron microscopy (TEM) (Figure 1d). For each mitochondrion, we measured the shortest outer-membrane-to-outer-membrane distance to its nearest neighbor. Blue light activation shifted the nearest-neighbor distance distribution toward shorter values (Figure 1e and S3; P < 0.0001). Using a TEM-defined contact threshold of <25 nm, the fraction of mitochondria participating in close contacts was also markedly increased following illumination (Figure 1f). Together, these measurements provide ultrastructural evidence that OptoMitoTether increases mitochondrial proximity.

To visualize the temporal relationship between CRY2PHR oligomerization and mitochondrial outer-membrane organization, we performed live-cell imaging using TOM20-EGFP together with CRY2PHR-mCherry-Miro1TM (Figure 1g). During 0–5 min of blue light exposure, CRY2PHR puncta progressively accumulated at interfaces between spatially distinguishable TOM20-positive mitochondrial profiles. These mitochondrial profiles remained morphologically distinguishable throughout the recorded interval, with no detectable morphological fusion in the tracked events. Thus, CRY2PHR oligomerization was temporally associated with increasing apposition of distinct mitochondrial outer-membrane profiles.

By comparing relative location of contacts (mCherry puncta) with respect to the morphology of mitochondria (labeled by MTDR), we found various types of mitochondrial contacts (Figure 1f, 1i), such as tip-to-tip, side-by-side, and tip-to-side configurations. Specifically, high-resolution SIM imaging well-resolved these distinct geometric contact modes at the single-organelle level: at site i, the tip of one mitochondrion was observed simultaneously interacting with the tips of two adjacent mitochondria. In contrast, lateral side-by-side alignments were found at sites ii and vi. Additionally, distinct tip-to-side junctions were clearly displayed at sites iii and iv, while a classical tip-to-tip docking between two mitochondrial ends was captured at site v.

To further characterize the precise temporal dynamics of this process, single-cell tracking was performed over a 30-minute period. This revealed a rapid accumulation of CRY2PHR puncta within 5 min of blue light exposure, followed by a gradual, time-dependent dissociation of these oligomers after shifting back to the dark (Figure S4). Although this short-term imaging primarily captured the reversible assembly of optogenetic components rather than the bulky organelles, a subsequent long-term study successfully confirmed the reversibility of mitochondrial morphology. Consistent with this rapid reversal, no persistent tethering was evident at 24 h; a 24-hour dark incubation allowed the mitochondria to disperse freely, fully recovering their native reticular structures and returning the quantified contact ratio back to the basal level (Figure S5a, S5b).

When half of a culture dish was covered with aluminum foil during blue light exposure; only the uncovered half resulted in enhanced mito-contacts, demonstrating spatial control (Figure S6). Meanwhile, cells transfected with CRY2PHR-mCherry-Miro1TM under darkness, or non-transfected cells under light illumination, do not show increased mito-contacts to non-transfected cells (Figures S7, S8). These results confirmed that both blue light and photoactivatable proteins are essential for enhancing mitochondrial interconnection. Furthermore, optogenetically induced mito-contacts were also observed in Michigan Cancer Foundation-7 (MCF-7) cells and neonatal human dermal fibroblasts (HDFn) (Figure S9). Together, these experiments demonstrate that OptoMitoTether provides reversible and spatially controlled induction of inter-mitochondrial tethering.

### OptoMitoTether elevates mitochondrial membrane potential without increasing ROS during acute activation

In this section, we sequentially subjected Control and OptoMitoTether cells to dark conditions, a 20 min blue light activation, and a 3 h damaging blue light exposure (all 300 μW/cm² light density) to track the cellular responses (Figure 2b). The 3h duration for the damaging exposure was determined based on a preliminary phototoxicity test, which revealed that blue light significantly compromised HeLa cell viability starting from 3 h of exposure (Figure S10). Meanwhile, the exposure within 1 h doesn’t cause detectable phototoxicity.

**Figure 2.**
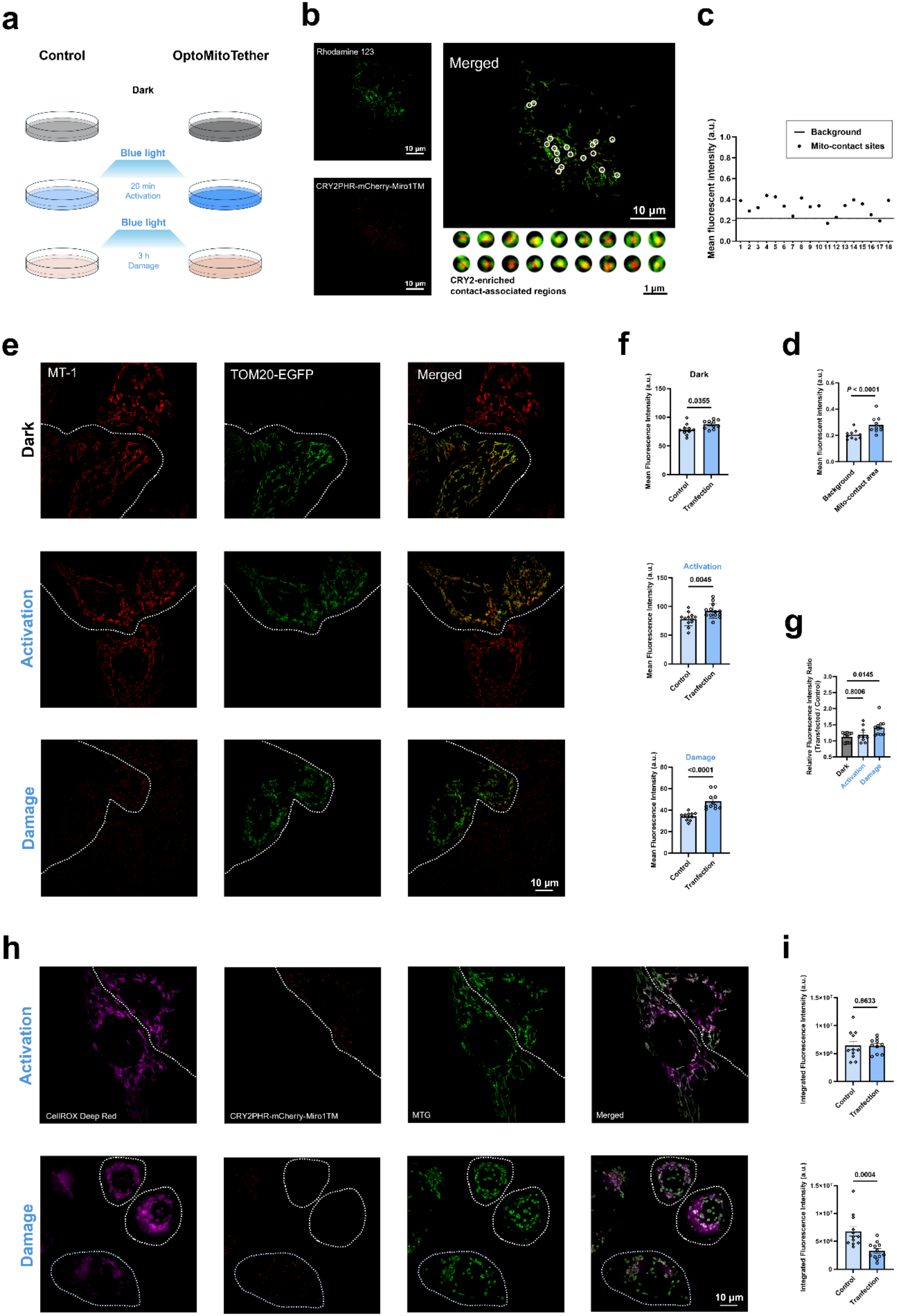
OptoMitoTether elevates mitochondrial membrane potential without increasing ROS during acute activation. **(a)** Experimental design. Control and OptoMitoTether-expressing HeLa cells were examined under dark conditions, following 20 min of blue light activation, or following 3 h of damaging blue light exposure. **(b)** Representative SIM images of Rh123-labeled OptoMitoTether-expressing HeLa cells following acute activation. Rh123 is shown in green and CRY2PHR-mCherry in red. Circles indicate CRY2PHR-enriched contact-associated regions used for local MMP analysis. **(c)** Rh123 fluorescence intensity for individual contact-associated regions relative to the mitochondrial background. **(d)** Mean Rh123 fluorescence intensity in mitochondrial background regions and CRY2PHR-enriched contact-associated regions. Data are mean ± SEM, (n = 10). P < 0.0001, two-tailed Student’s t test.; n = 11. P < 0.0001, two-tailed Student’s t test. **(e)** Representative images from the complementary MT-1 assay under dark, 20 min activation, and 3 h damage conditions. MT-1/mCherry-channel fluorescence is shown in red and TOM20-EGFP in green. Dashed outlines distinguish the transfected cells from the control (untransfected) group based on the presence or absence of the TOM20-EGFP signal. **(f)** Mean red-channel fluorescence intensity in control and OptoMitoTether-expressing cells under each condition. Dark, P = 0.0355; 20 min activation, P = 0.0045; 3 h damage, P < 0.0001. **(g)** Ratio of mean red-channel fluorescence intensity in OptoMitoTether-expressing cells relative to control cells across the three conditions. The ratio didn’t differ between dark and acute activation (P = 0.8006) but was significantly greater under prolonged damaging blue light exposure in the indicated comparison (P = 0.0145). Data were analyzed using one-way ANOVA followed by Tukey’s test. **(h)** Representative images of CellROX Deep Red (ROS Probe) fluorescence following acute activation and prolonged damaging blue light exposure. CellROX is shown in magenta, CRY2PHR-mCherry in red, MTG mitochondrial signal in green, and merged images at right. **(i)** Quantification of integrated CellROX fluorescence. Under acute activation, ROS levels didn’t differ between control and OptoMitoTether-expressing cells (P = 0.8633). Following 3 h of damaging blue light exposure, ROS levels were significantly lower in OptoMitoTether-expressing cells (P = 0.0004). Each point represents one cell; two-tailed Student’s t tests.

To figure out whether OptoMitoTether induced tethering was associated with changes in MMP. Rhodamine 123 (Rh123) was first used to quantify MMP in mitochondrial regions surrounding CRY2PHR-enriched puncta following acute photoactivation (Figure 2b–2d). CRY2PHR puncta were used as spatial landmarks to define contact-associated regions, and Rh123 fluorescence within mitochondrial pixels surrounding these puncta was compared with fluorescence in the remaining mitochondrial regions. Most selected regions in Figure 2b show higher Rh123 fluorescence than background baseline (Figure 2c) and the average intensity of contact is significantly higher than the background mitochondrial region (Figure 2d) supporting a local association between mitochondrial tethering and elevated MMP.

We next used MT-1 as a complementary cell-level MMP-associated indicator under three conditions: dark, 20 min blue light activation, and 3 h damaging blue light exposure (Figure 2e–2g). Because MT-1 and mCherry were detected within the same red acquisition channel, a modest transfection-associated fluorescence offset was present even under dark conditions. We therefore compared the transfected-to-control red-channel fluorescence intensity ratio across experimental conditions, rather than relying solely on the absolute fluorescence intensity difference between transfected and control cells. The ratio didn’t differ between dark and acute activation conditions (P = 0.8006), whereas it was significantly larger under prolonged damaging blue light exposure (P = 0.0145 for the indicated comparison; Figure 2g). Thus, the pronounced MT-1-channel difference observed during prolonged stress exceeds the baseline transfection-associated fluorescence offset.

Because increased MMP can under some conditions be accompanied by increased oxidative stress (Korshunov et al., 1997; Murphy, 2009), we next quantified ROS using CellROX DeepRed (Figure 2h, 2i). Following acute 20 min activation, ROS levels didn’t differ between control and OptoMitoTether-expressing cells (P = 0.8633). In contrast, after 3 h of damaging blue light exposure, ROS levels were significantly lower in OptoMitoTether-expressing cells than in controls (P = 0.0004). Thus, acute OptoMitoTether activation was not accompanied by a detectable increase in ROS, whereas during prolonged blue light stress OptoMitoTether expression was associated with reduced oxidative burden.

### OptoMitoTether preserves mitochondrial function during prolonged blue light stress

Having established that OptoMitoTether activation increases mitochondrial membrane-potential and reduces oxidative burden during prolonged blue light exposure, we next examined whether these differences were accompanied by preservation of mitochondrial function. Cells were divided into Control and Transfected (OptoMitoTether-expressing) groups and maintained either in darkness or under continuous blue light for 3 h (Figure 3a). Prolonged blue light exposure induced pronounced mitochondrial fragmentation in control cells, including an increased proportion of rounded mitochondria and reduced mitochondrial apposition (Figures 3b and S11). Under the same damaging condition, mitochondria in OptoMitoTether-expressing cells still exhibited fragmentation but showed a substantially higher frequency of contacts among mitochondrial fragments (Figure 3c). We therefore describe this organization as a contact-connected mitochondrial network rather than restoration of a fused tubular network.

**Figure 3.**
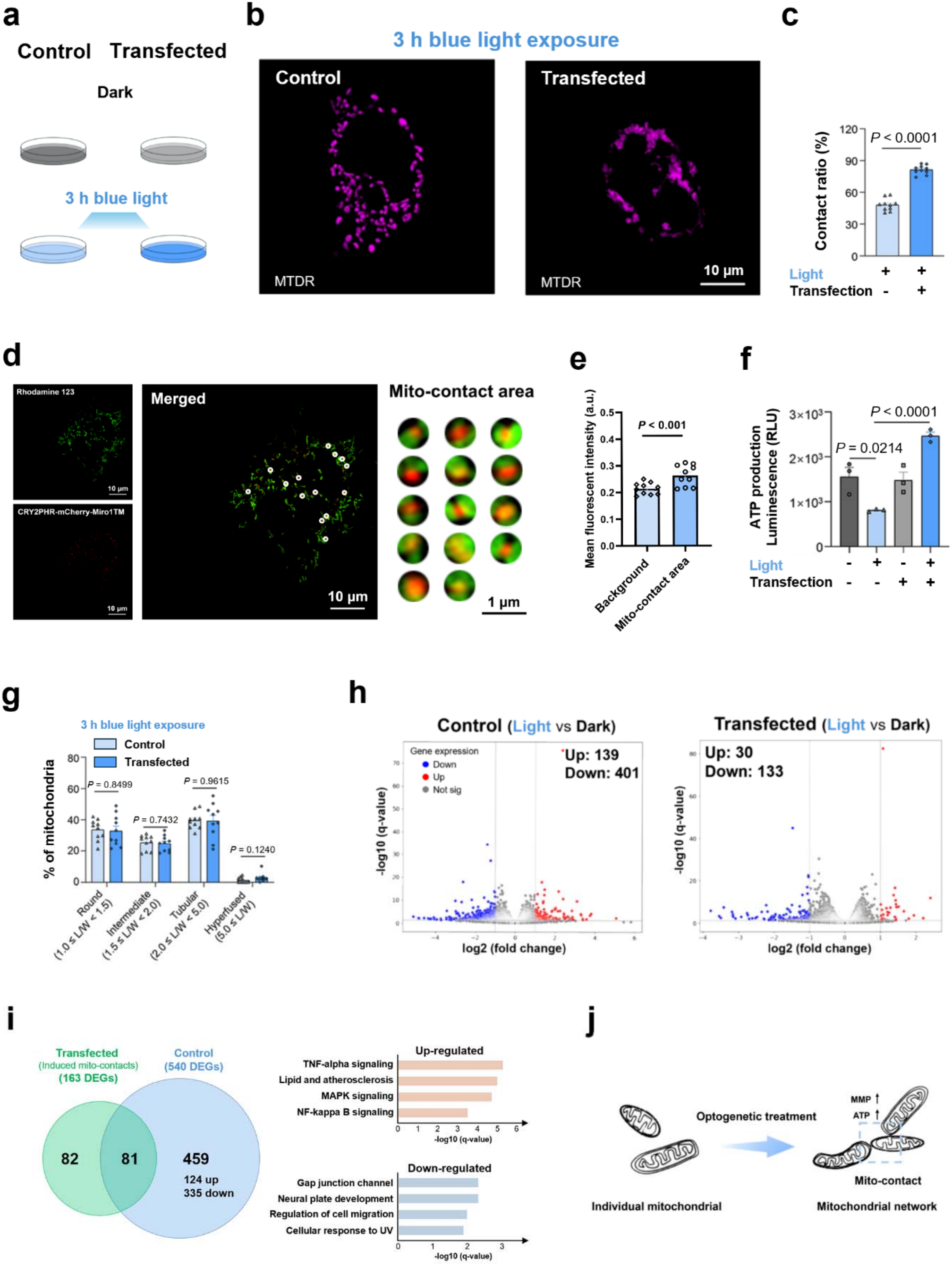
OptoMitoTether preserves mitochondrial function during prolonged blue light stress. **(a)** Schematic representation of the experimental conditions used to examine the effects of prolonged blue light exposure in control and OptoMitoTether-expressing HeLa cells. **(b)** Representative SIM images of control and OptoMitoTether-expressing HeLa cells following 3 h of blue light exposure (300 μW/cm²). Mitochondria were labeled with MTDR. **(c)** Mitochondrial contact ratio in control and OptoMitoTether-expressing cells following 3 h of blue light exposure. Data are presented as mean ± SEM (n = 10 cells from three independent experiments). P < 0.0001, two-tailed Student’s t test. **(d)** Representative SIM images of Rh123 labeled OptoMitoTether-expressing HeLa cells following 3 h ofblue light exposure. Rh123 is shown in green and CRY2PHR-mCherry in red. Enlarged regions showCRY2PHR-enriched contact-associated mitochondrial regions. **(e)** Mean Rh123 fluorescence intensity in mitochondrial background regions and CRY2PHR-enrichedcontact-associated regions following 3 h of blue light exposure. Data are presented as mean ± SEM (n = 10cells). P < 0.001, two-tailed Student’s t test. **(f)** The ATP production of HeLa cells with different treatments. mean ± SEM (n = 3 independent samples mean ± SEM (n = 3 independent samples). **(g)** Quantitative analysis of mitochondrial morphology in (b). Mitochondria were classified as hyperfused, tubular, intermediate, or round according to aspect ratio. Data are presented as mean ± SEM (n = 10 cells). **(h)** The differential gene expression between dark and 3h light treated cells from RNA-sequence data of all four groups HeLa cells. The left one was Control groups (light vs dark), mapping the 139 upregulated genes (red) and 401 downregulated genes (blue); the right one was Transfected groups (light vs dark), mapping the 30 upregulated genes (red) and 133 downregulated genes (blue). **(i)** Overlap between differentially expressed genes identified in control and OptoMitoTether-expressing cells and functional annotation of blue light-specific transcriptional changes. Of the 540 differentially expressed genes identified in control cells, 459 were unique to the control condition. **(j)** Schematic summary of the proposed functional consequences of OptoMitoTether-mediated mitochondrial tethering during prolonged blue light stress. Statistical differences between the experimental groups were analyzed using a double-tailed Student’s t test. All P values less than 0.05 were considered to indicate statistical significance.

Long-time blue LED light exposure causes mitochondrial depolarization (Hao et al., 2023), but mito-contacts can maintain relatively high MMP levels at the mito-contact area (Figures 3d, 3e) as well as the relative high whole cell level MMP shown as the damage condition in Figures 2e, 2f, consistent with prior findings that close apposition of charged membranes or proteins can markedly enhance membrane potential (Bu et al., 2016; Gurunian et al., 2025). Damaged cells, particularly those with impaired mitochondria, produce less energy yet require more energy for repair than under normal conditions (Flood et al., 2023; Milanese et al., 2019). Since MMP drives ATP synthesis by providing the electrochemical proton gradient (Duchen, 2000; Romero-Garcia and Prado-Garcia, 2019), we quantified ATP production to assess whether locally high MMP would benefit mitochondrial function. As shown in Figure 3f, blue light exposure significantly reduced ATP levels in control cells, whereas OptoMitoTether-expressing cells maintained substantially higher ATP levels under the same condition. ATP levels in the light-treated OptoMitoTether group exceeded those measured in the untreated control condition. The mechanism underlying this increase in ATP production is not established by the present data; we therefore interpret this observation conservatively as evidence that OptoMitoTether-expressing cells maintain high energetic capacity during prolonged blue light stress.

As mitochondrial fusion is known to support mitochondrial function (Guo et al., 2023), we conducted a detailed analysis of mitochondrial morphology using SIM imaging. Morphological analysis showed that the protective phenotype was not accompanied by restoration of a uniformly hyperfused mitochondrial morphology (Figure 3g). Instead, the system linked fragmented mitochondria into an extensive network, thereby increasing the number of contact sites among them (Figure 3c).

We then performed RNA sequencing to determine whether OptoMitoTether activation induces changes in gene transcription. Following 3 h of blue light exposure, control cells showed 540 differentially expressed genes (DEGs) (401 downregulated and 139 upregulated), whereas OptoMitoTether-expressing cells showed 163 DEGs (133 downregulated and 30 upregulated; Figure 3h). Among the 540 DEGs regulated by blue light, there were 459 unique to the non-transfected group (335 downregulated and 124 upregulated; Figure 3i). The blue light-specific upregulated DEGs were mainly associated with inflammatory signaling pathways (e.g., TNF-α, MAPK, and NF-κB signaling) and lipid and atherosclerosis signaling. In contrast, the downregulated DEGs are linked to gap junction channels, neural plate development, cell migration, and cellular responses to UV exposure. These results are associated with an attenuated transcriptional response involving inflammatory signaling and cellular- remodeling pathways.

Several mitochondria-associated genes showed differential expression in control cells, including PET100, SDHAF4, and HOGA1. PET100 is involved in complex IV assembly (Church et al., 1996; Vidoni et al., 2017), SDHAF4 is associated with complex II assembly, and HOGA1 participates in mitochondrial metabolism (Cochat and Rumsby, 2013; Li et al., 2023; Vidoni et al., 2017). These individual gene-level findings are interpreted descriptively rather than as validated molecular mediators. At the gene-set level, mitochondrial GO and KEGG enrichment analyses revealed significant blue light-associated changes in control cells but no significant terms in OptoMitoTether-expressing cells at the same statistical threshold (Figure S12). These transcriptional patterns are consistent with an attenuation of the blue light-associated transcriptional stress response but do not establish the molecular mechanism underlying the ATP phenotype.

### OptoMitoTether preserves mitochondrial function in human retinal cells during prolonged blue light exposure

Considering that the eyes and skin are commonly exposed to natural or artificial blue light throughout the day, the exposure duration for the damage model was extended from hours to a full day (Cougnard-Gregoire et al., 2023; Nash et al., 2019). We applied the optogenetic mito-contact system to severely damaged human retinal cells (ARPE-19) caused by prolonged low-intensity blue light exposure (Chakravarthy et al., 2024). After 24 hours of blue light exposure, the viability of ARPE-19 cells drops below 50%, underscoring the substantial cytotoxicity induced by prolonged blue light exposure (Figure S13). As shown in Figure 4a, long time exposure induced morphological changes in mitochondria within human retinal cells, transforming them from elongated shapes into fragmented, granular forms-hallmarks of mitochondrial damage. Although mitochondria remained extensively fragmented following prolonged blue light exposure, OptoMitoTether-expressing cells showed substantially greater physical association among mitochondrial fragments (Figure 4b), producing a contact-connected organization rather than restoration of a fused tubular network.

**Figure 4.**
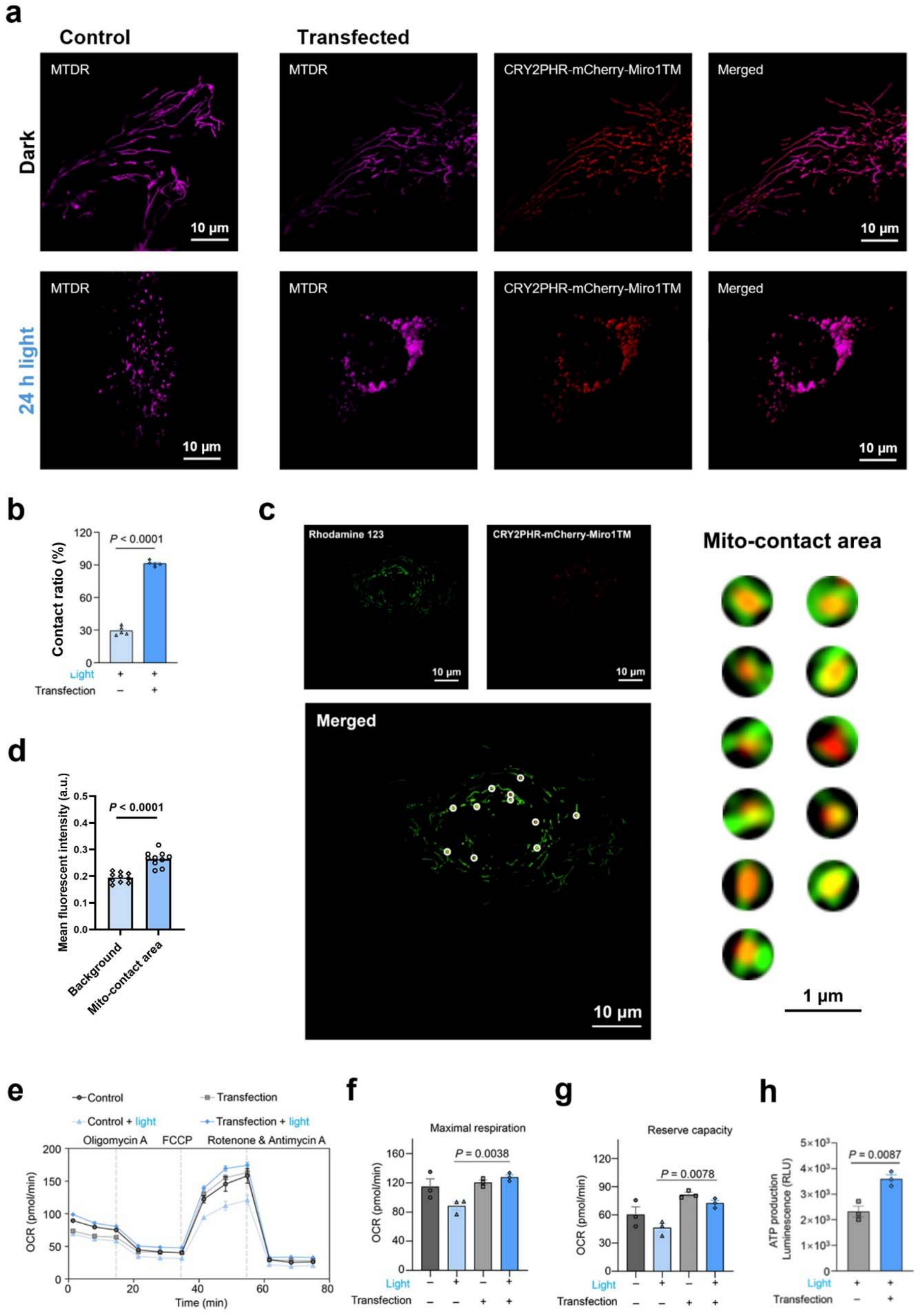
OptoMitoTether preserves mitochondrial bioenergetic function in human retinal cells during prolonged blue light exposure. **(a)** Representative SIM images of ARPE-19 cells under dark conditions or following 24 h of blue light exposure (300 μW/cm²), with or without OptoMitoTether expression. Mitochondria were labeled with MTDR. Prolonged illumination caused extensive mitochondrial fragmentation, whereas OptoMitoTether-expressing cells showed increased physical association among mitochondrial fragments. **(b)** Mitochondrial contact ratio in ARPE-19 cells following prolonged blue light exposure. Data are presented as mean ± SEM (n = 5 cells). P < 0.0001, two-tailed Student’s t test. **(c)** Representative SIM images of Rh123-labeled OptoMitoTether-expressing ARPE-19 cells following 24 h of blue light exposure. Rh123 is shown in green and CRY2PHR-mCherry in red. Enlarged regions show CRY2PHR enriched contact-associated mitochondrial regions. **(d)** Mean Rh123 fluorescence intensity in mitochondrial background regions and CRY2PHR enriched contact-associated regions. Contact-associated regions showed 36.5% greater fluorescence than the mitochondrial background. Data are presented as mean ± SEM (n = 10 cells). P < 0.0001, two-tailed Student’s t test. **(e)** Oxygen-consumption-rate (OCR) profiles from the mitochondrial stress test in control and OptoMitoTether-expressing ARPE-19 cells under dark and prolonged blue light conditions. Oligomycin A, FCCP, and rotenone/antimycin A were sequentially applied as indicated. Data are presented as mean ± SEM (n = 3 independent samples). **(f)** Maximal mitochondrial respiration measured from the OCR stress test. Data are presented as mean ± SEM (n = 3 independent samples). Statistical comparisons are indicated in the figure. **(g)** Mitochondrial reserve respiratory capacity measured from the OCR stress test. Data are presented as mean ± SEM (n = 3 independent samples). Statistical comparisons are indicated in the figure. **(h)** ATP levels in control and OptoMitoTether-expressing ARPE-19 cells following 24 h of blue light exposure. Data are presented as mean ± SEM (n = 3 independent samples). P = 0.0087, two-tailed Student’s t test. Statistical differences between the experimental groups were analyzed using a double-tailed Student’s t test. All P values less than 0.05 were considered to indicate statistical significance.

MMP analysis shows that the mito-contact area consistently exhibited significantly higher fluorescence compared with the background (Figure 4c). Specifically, the fluorescence intensity at mito-contacts was 36.5% higher than the background average (Figure 4d). Functional analysis using the Seahorse assay confirmed that blue light exposure impairs mitochondrial function, whereas light-induced mito-contacts help preserve it (Figures 4e–4g). ATP quantification (Figure 4h) ATP quantification further showed that OptoMitoTether-expressing ARPE-19 cells maintained higher ATP levels than control cells under the same prolonged blue light condition. These findings suggest that the optogenetic mito-contact system can safeguard mitochondrial function following severe light-induced damage in vitro.

### OptoMitoTether extends lifespan of *C. elegans* under chronic blue light exposure

To determine whether OptoMitoTether-mediated mitochondrial tethering is also effective in vivo, we expressed the system in *C. elegans* (Figure 5a). Prolonged blue light exposure produced pronounced mitochondrial fragmentation in control animals (Figure 5b), a hallmark of mitochondrial dysfunction (Johnson et al., 2021; Wai et al., 2015), while OptoMitoTether-expressing animals showed a more contact-associated mitochondrial organization under the same condition (Figure 5c), demonstrating that the optogenetic mito-contact system is functional and effective *in vivo*.

**Figure 5.**
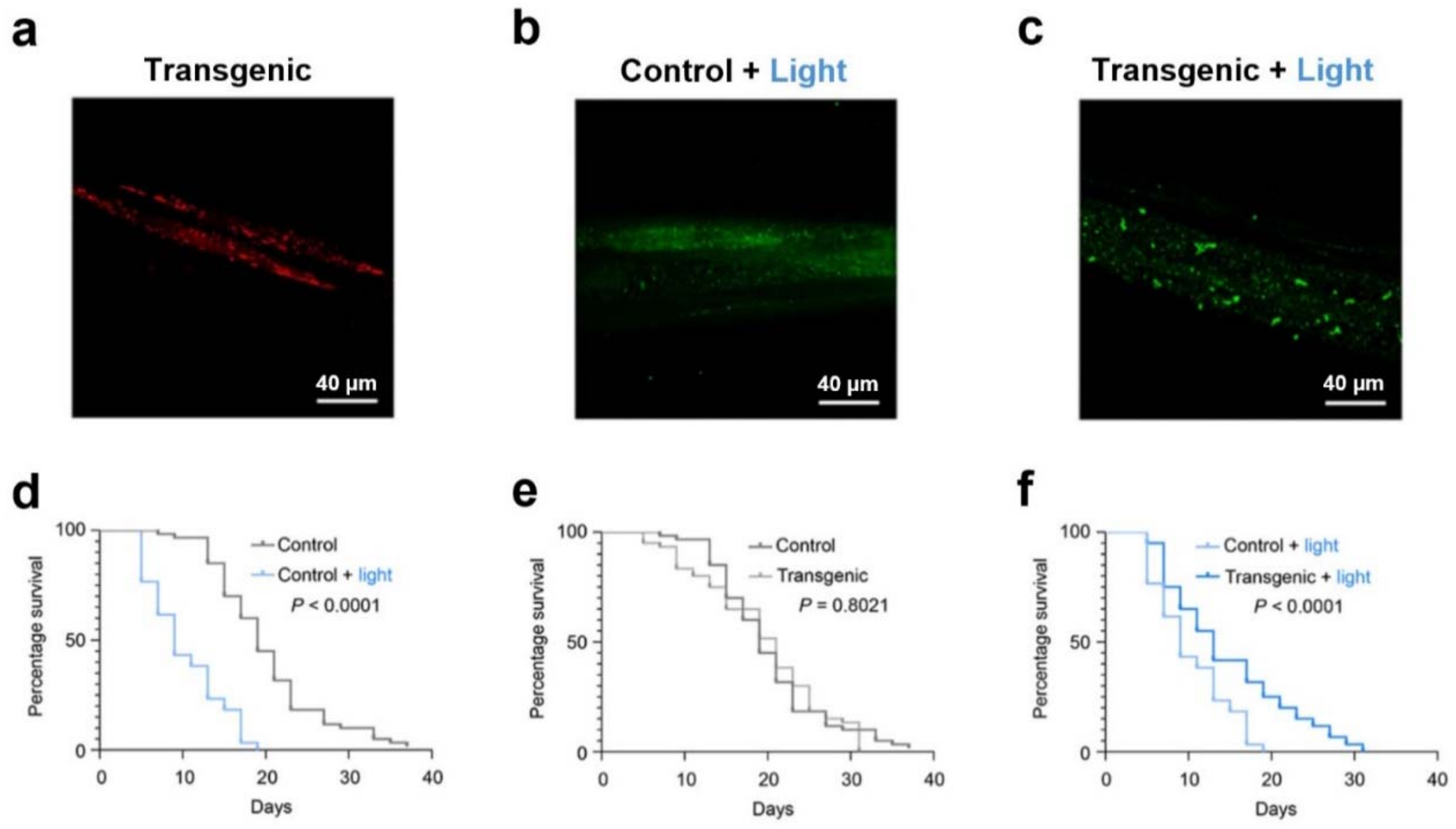
OptoMitoTether extends *C. elegans* lifespan during chronic blue light exposure. **(a)** The fluorescent image of *C. elegans* animal carrying the CRY2PHR-mCherry-Miro1TM transgenic array. **(b)** Representative fluorescence image of mitochondrial organization in a control *C. elegans* animal following chronic blue light exposure. Mitochondria were labeled with MTG. **(c)** Representative fluorescence image of mitochondrial organization in an OptoMitoTether-expressing *C.elegans* animal during chronic blue light exposure. Mitochondria were labeled with MTG.**(d)** Survival curves of control animals maintained in darkness or under chronic blue light exposure (n = 60 animals per group). Chronic blue light exposure significantly shortened lifespan (P < 0.0001, log-rank Mantel–Cox test). **(e)** Survival curves of non-transgenic control and OptoMitoTether-transgenic animals maintained without blue light exposure (n = 60 animals per group). No significant lifespan difference was observed between groups (P = 0.8021, log-rank Mantel–Cox test). **(f)** Survival curves of control and OptoMitoTether-transgenic animals during chronic blue light exposure (n =60 animals per group). OptoMitoTether expression significantly increased lifespan under chronic blue light exposure (P < 0.0001, log-rank Mantel–Cox test).

Lifespan is intricately tied to the maintenance of mitochondrial quality control (Sharma et al., 2019). Building on this finding, we subsequently investigated the effects of mito-contacts on the lifespan of *C. elegans* under long-term blue light exposure. The constant blue light treatment was initiated on Day 1 of adulthood and continued until death. Compared with the untreated control group, animals exposed to blue light exhibited significantly reduced lifespans (Figure 5d). Importantly, no significant differences were observed between the untreated control and untreated transgenic worms (Figure 5e). However, transgenic worms subjected to blue light treatment showed a significant increase in lifespan compared with the non-transfected control group under the same blue light condition (Figure 5f). These results indicate that mito-contact induced by OptoMitoTether counteracts the detrimental effects of long-term blue light exposure, thereby contributing to lifespan extension in damaged organisms.

## Discussion

In this study, we developed OptoMitoTether, an optogenetic system that enables reversible manipulation of inter-mitochondrial proximity with high spatial and temporal control. Blue light activation increased the proportion of mitochondria participating in close appositions, shortened mitochondrial nearest-neighbor distances at the ultrastructural level, and promoted the formation of CRY2PHR-enriched interfaces between distinct mitochondrial outer-membrane profiles. These structural changes were accompanied by elevated membrane potential in contact-associated mitochondrial regions. Under prolonged blue light stress, OptoMitoTether-expressing cells further showed reduced ROS accumulation, preserved ATP production, and an attenuated transcriptional response to mitochondrial stress. Importantly, the protective phenotype extended beyond HeLa cells: OptoMitoTether preserved mitochondrial respiratory function and ATP production in human retinal cells and increased the lifespan of C. elegans during chronic blue light exposure. Together, these results identify controlled inter-mitochondrial tethering as a means of modulating mitochondrial organization and bioenergetic resilience without directly targeting the canonical fusion–fission machinery.

A central implication of these findings is that mitochondrial organization may influence bioenergetic function at a level distinct from complete organelle fusion. Mitochondria are highly dynamic, and fusion and fission are well-established mechanisms for redistributing mitochondrial contents, maintaining mitochondrial quality, and adapting network architecture to cellular demands (Chen and Chan, 2009; Detmer and Chan, 2007; Duchen, 2000; McBride et al., 2006; Nunnari and Suomalainen, 2012; Szabadkai et al., 2004). Dysregulation of these processes is closely linked to mitochondrial dysfunction in aging, neurodegeneration, and metabolic and cardiovascular disease (Chen et al., 2007; Dorn, 2015; Pokharel et al., 2024; Shah et al., 2019; Suárez-Rivero et al., 2016; Sun et al., 2016; Szendroedi et al., 2011; Wang et al., 2009; Yapa et al., 2021). Our results suggest that close physical association between otherwise distinguishable mitochondria may provide an additional mode of organization. Even after prolonged blue light exposure, when mitochondria remained largely fragmented, OptoMitoTether increased the frequency of physical contacts among these fragments rather than restoring a uniformly elongated or hyperfused network. This contact-connected organization was nevertheless associated with higher MMP and ATP levels under stress. Thus, functional coordination within a mitochondrial population may not require all participating mitochondria to merge into a continuous organelle.

How mitochondrial proximity influences membrane potential remains an important mechanistic question. MMP represents the electrochemical proton gradient across the inner mitochondrial membrane and is central to ATP synthesis, protein import, ion transport, and other mitochondrial functions (Miller and Sheetz, 2004; Novak, 2012; Paschen and Neupert, 2001; Sato et al., 2019; Scott et al., 1980; Youle and Narendra, 2010; L D Zorova et al., 2018; Ljubava D Zorova et al., 2018). Previous work has suggested that close membrane apposition can alter local electrostatic conditions and membrane potential (Bu et al., 2016), providing a potential physical basis for the local MMP enhancement observed around OptoMitoTether-associated contact regions. In addition, naturally occurring inter-mitochondrial junctions have been proposed to facilitate structural and functional coordination among neighboring mitochondria. Picard and colleagues showed that mitochondria at such junctions can exhibit coordinated cristae organization and that experimentally enforced mitochondrial proximity can promote trans-mitochondrial structural coordination (Picard et al., 2015). This framework is particularly relevant to our observations because OptoMitoTether generates reproducible geometries of mitochondrial appositions, including tip-to-tip, tip-to-side, and side-by-side arrangements. At these contact sites, neighboring mitochondria often adopt similar overall orientations, suggesting that external mitochondrial geometry can influence the organization of internal structures. As shown in Figure 1g, tip-to-tip and tip-to-side configurations favor a serial, tandem alignment of cristae across the contact interface (resembling a continuous staircase), whereas lateral side-by-side configurations support parallel, ladder-like cristae registries. Although individual cristae cannot be resolved by our SIM imaging—preventing direct verification of their alignment—the observed orientation of adjacent mitochondrial profiles is consistent with the hypothesis that physical apposition organizes mitochondria across multiple structural scales to facilitate bioenergetic coupling.

The distinction between tethering and fusion is also important for understanding how OptoMitoTether reorganizes mitochondria. Mitochondrial fusion requires tightly regulated membrane approximation and remodeling, involving dedicated fusion machinery on both the outer and inner mitochondrial membranes (Chen and Chan, 2009; Detmer and Chan, 2007; Guo et al., 2023; Nunnari and Suomalainen, 2012). Studies of membrane fusion further indicate that opposing membranes must reach very short separations before fusion-pore formation can proceed (Bu et al., 2016). In the time-lapse TOM20-EGFP recordings, mitochondrial profiles associated with newly formed CRY2PHR puncta remained morphologically distinguishable over the observation period, and prolonged blue light exposure produced numerous contact-connected mitochondrial fragments rather than widespread fusion. One possible explanation is that membrane-anchored CRY2PHR oligomers themselves occupy the space between adjacent outer mitochondrial membranes. The CRY2PHR domain is a relatively large protein module, and oligomerization of membrane-anchored CRY2PHR molecules could impose steric constraints that stabilize close apposition while limiting the very short membrane separations required for merger (Palayam et al., 2021). Such a configuration would allow mitochondria to remain closely tethered while retaining their individual membrane identities. This interpretation is consistent with our observation that OptoMitoTether readily promotes mitochondrial tethering while detectable fusion remains uncommon. At the same time, the present imaging doesn’t exclude transient fusion events that are not morphologically resolved, so our data supports tethering as a stable and functionally relevant mitochondrial state without requiring widespread fusion.

The functional consequences of this tethered state became particularly apparent under mitochondrial stress. Short-term OptoMitoTether activation increased MMP locally around CRY2PHR-enriched contact-associated regions without producing a detectable increase in ROS. This distinction is important because mitochondrial hyperpolarization is not necessarily beneficial: excessively high proton motive force can enhance mitochondrial ROS production (Cunha et al., 2011; Korshunov et al., 1997). Conversely, mitochondrial depolarization compromises ATP production and multiple homeostatic processes (Glancy and Balaban, 2012; Green and Reed, 1999; Hüttemann et al., 2008; Ly et al., 2003; Rossignol et al., 2004; Rovini et al., 2021; Santo-Domingo and Demaurex, 2010). Under the conditions examined here, enhanced contact-associated MMP was not accompanied by an acute oxidative penalty. During prolonged blue light exposure, the difference between OptoMitoTether-expressing and control cells became more pronounced at the cellular level, while ROS levels were reduced in OptoMitoTether-expressing cells. These findings suggest that mitochondrial tethering helps maintain a more favorable bioenergetic state when mitochondrial function is challenged rather than simply inducing indiscriminate hyperpolarization.

This protective effect is particularly relevant in the context of blue light stress. Prolonged blue light exposure has been reported to induce mitochondrial fragmentation, ROS production, mitochondrial DNA damage, MMP loss, and impaired mitochondrial function in multiple cell types, including retinal cells (Godley et al., 2005; Guo et al., 2020; Hao et al., 2023; Li et al., 2018; Tao et al., 2019; Wang et al., 2023; Yan et al., 2025). Consistent with these studies, prolonged illumination in our experiments reduced cell viability, disrupted mitochondrial morphology, and lowered ATP production in control cells. In contrast, OptoMitoTether-expressing cells maintained higher ATP levels and lower ROS under the same conditions. Because ATP production depends strongly on the proton-motive force generated by the respiratory chain (Hüttemann et al., 2008; L D Zorova et al., 2018; Ljubava D Zorova et al., 2018), preservation of MMP provides a plausible link between mitochondrial tethering and maintained energetic capacity, although the present data do not establish a single causal pathway from local MMP elevation to whole-cell ATP production.

The RNA-seq results provide an additional view of this stress-protective state. Blue light exposure produced substantially more transcriptional changes in control cells than in OptoMitoTether-expressing cells, including genes and pathways associated with inflammatory signaling, cellular remodeling, and mitochondrial function. Several mitochondria-associated genes involved in respiratory-chain assembly or metabolism, including PET100, SDHAF4, and HOGA1, were altered in control cells but not significantly changed in OptoMitoTether-expressing cells. PET100 is involved in cytochrome-c oxidase assembly (Church et al., 1996; Vidoni et al., 2017), whereas SDHAF4 contributes to complex II assembly and HOGA1 participates in mitochondrial metabolism. At the gene-set level, significant mitochondrial GO and KEGG changes were detected in control cells but not in OptoMitoTether-expressing cells. These transcriptional differences do not identify the molecular mechanism underlying the energetic phenotype, but they are consistent with a broader attenuation of the cellular response to prolonged mitochondrial stress.

The findings in ARPE-19 cells further extend the relationship between mitochondrial tethering and bioenergetic resilience. Retinal cells are especially relevant to blue light injury because visible-light exposure can directly affect mitochondrial integrity and has been implicated in retinal phototoxicity (Chakravarthy et al., 2024; Li et al., 2018; Tao et al., 2019; Wang et al., 2023; Yan et al., 2025). After prolonged illumination, control ARPE-19 cells showed marked mitochondrial fragmentation and impaired respiratory function. OptoMitoTether didn’t restore a conventional fused tubular morphology; instead, fragmented mitochondria became more extensively contact-connected. This structural state was associated with higher contact-associated MMP, greater maximal respiratory capacity, greater reserve capacity, and higher ATP production. The convergence of these measures indicates that the protective effect of OptoMitoTether extends beyond a single fluorescence-based MMP readout to mitochondrial respiratory performance under severe stress.

The *C. elegans* experiments further demonstrated that the consequences of mitochondrial tethering can extend to the organismal level. Mitochondrial quality control and dynamics are closely linked to organismal aging and longevity (Johnson et al., 2021; Sharma et al., 2019; Wai et al., 2015). Chronic blue light exposure has also been shown to shorten lifespan and induce neurodegenerative phenotypes in model organisms (Nash et al., 2019). Consistent with these observations, chronic blue light exposure shortened lifespan in control animals in our study. In contrast, animals expressing OptoMitoTether survived substantially longer under the same illumination, whereas expression of the system in the absence of blue light activation didn’t significantly alter lifespan. This pattern suggests that the benefit of OptoMitoTether emerges specifically when the optogenetic system is engaged in a damaging light environment rather than from constitutive expression of the construct itself. This stimulus dependence is a distinctive feature of the system. In the present paradigm, blue light serves different functions over different time scales: brief illumination activates CRY2PHR oligomerization and mitochondrial tethering (Duan et al., 2015; Zhang and Cui, 2015), whereas prolonged exposure produces mitochondrial stress (Godley et al., 2005; Hao et al., 2023; Li et al., 2018; Tao et al., 2019; Wang et al., 2023; Yan et al., 2025). OptoMitoTether therefore creates a stimulus-coupled form of mitochondrial regulation in which the same environmental input that eventually imposes stress also recruits a compensatory structural response. This differs from constitutive manipulation of mitochondrial fusion, fission, or metabolism and allows mitochondrial organization to be controlled with temporal and spatial precision. Such optogenetic control of organelle interactions has increasingly been used to dissect functional relationships between mitochondria and other organelles (Bai et al., 2025; Qiu et al., 2025, 2023, 2022), and OptoMitoTether extends this strategy specifically to reversible mitochondria–mitochondria interactions.

More broadly, these findings suggest that mitochondrial networks should not be described solely along a fused-versus-fragmented axis. In cardiac and other highly energetic tissues, physically clustered mitochondria can display coordinated electrical and metabolic behavior even when individual mitochondrial boundaries remain distinguishable (Amchenkova et al., 1988; Kurz et al., 2017; Sun et al., 2020). Our results support a related view in which the spatial arrangement and contact architecture among individual mitochondria represent an additional dimension of mitochondrial organization. By allowing mitochondrial proximity to be manipulated without directly perturbing the canonical fusion– fission machinery, OptoMitoTether provides a way to separate the functional consequences of physical apposition from those of complete membrane fusion.

In conclusion, OptoMitoTether enables reversible control of inter-mitochondrial proximity and reveals a functional relationship between mitochondrial tethering, membrane potential, and bioenergetic resilience. By increasing close mitochondrial apposition, the system preserves mitochondrial function during prolonged blue light stress in cultured cells, human retinal cells, and C. elegans. These findings establish inter-mitochondrial tethering as a distinct dimension of mitochondrial organization and provide a framework for investigating how physical mitochondrial interactions contribute to cellular energy homeostasis.

## Author Contributions

H.B., A.L.Z., K.Z., and J.D. conceived the project. Y.W. and K.Q. carried out cell imaging experiments and analysis, ATP experiment and RNA-seq. W.Z. did the Seahorse experiment. P.A. performed the lifespans and imaging of animals. T.H.G. injected and screened transgenic animals. E.L. did procedures for contact determination. Z.T. assisted with data collection. X.X. drew the schematic. P.K. analyzed RNA-seq data. Y.W., K.Q. and J.D. wrote the manuscript. T.H., N.-P.T., D.S., H.B., A.L.Z., and K.Z. revised the manuscript. All authors participated in discussions on results and in preparing the manuscript.

## Notes

The authors declare no competing financial interest.

## ACKNOWLEDGMENTS

This work was supported by the National Institutes of Health (NIH R35GM128837 to J.D.; R01GM132438, R01MH124827, and National Science Foundation Science and Technology Center for Quantitative Cell Biology, 2243257 to K.Z.; R01AG058741 and R01AG075156 to H.B.), the National Science Foundation (NSF CAREER 2046984 to H.B.), and the Hevolution Foundation (HF-GRO-23-1199062-14). Additional support was provided by the Cincinnati Children’s Research Foundation in the form of a Trustee Award to A.L.Z. We acknowledge the support of the Cincinnati Children’s Confocal Imaging Core SCR_022628. Caenorhabditis elegans strain N2 was obtained from the Caenorhabitis Genetics Center project funded by NIH Office of Research Infrastructure Programs (P40 OD010440).

## Methods

### Materials

MitoTracker^TM^ Green FM (MTG, #M7514) and MitoTracker^TM^ Deep Red FM (MTDR, #M22426) were obtained from Invitrogen (Thermo Fisher Scientific, USA). Penicillin–streptomycin (#15140163, 10,000 units/mL), fetal bovine serum (FBS, #26140079), DMEM (#11965092), DMEM/F-12 (#11320082) and geneticinTM selective antibiotic (#10131035) were all obtained from Gibco (Thermo Fisher Scientific, USA). Phosphate-buffered saline (PBS, #SH30256.01) was sourced from Hyclone (GE Healthcare Life Sciences, USA). CRY2PHR-mCherry-Miro1TM (Addgene plasmid, #102247) was a gift from Bianxiao Cui. TurboFect^TM^ Transfection Reagent (#R0532), CellROX™ Deep Red Reagent (#C10422), Invitrogen™ ATP Determination Kit (#A22066) and Rhodamine 123 (#R302) were purchased from Thermo Scientific (Thermo Fisher Scientific, USA). Cell Counting Kit-8 (CCK-8) and MT-1 MitoMP Detection Kit (MT13-10) were purchased from Dojindo (Dojindo Molecular Technologies, Inc., Japan).

### Cell Culture and Transfection

The HeLa cell line, MCF-7 cell line, and HDFn cell line were cultured in DMEM supplemented with 10% FBS and 100 units/mL of penicillin-streptomycin. ARPE-19 cell line (arising retinal pigment epithelia, #CRL-2302, ATCC), were cultured in DMEM/F-12 supplemented with 10% FBS and 100 units/mL of penicillin-streptomycin. The cell cultures were maintained at 37 °C in a 5% CO2 cell incubator (Thermo Fisher Scientific, USA) with 100% humidity. Transfection was carried out utilizing TurboFectTM Transfection Reagent following the instructions provided by the manufacturer. We suggest using 2 μg CRY2PHR-mCherry-Miro1TM with 6 μL of TurboFect for 35 mm dish. The mixtures of DNA and transfection reagent were added dropwise to the cell cultures and incubated for 6 hours. After the incubation period, cell cultures were replenished with complete culture medium. To select the expressed cells, geneticin^TM^ selective antibiotic at 500 ng/mL was treated with transfected cells.

### Cell Imaging

Nikon Structured Illumination Microscopy (N-SIM, Nikon Corporation, Japan), a super-resolution microscope system was applied for live cell imaging in this study. The microscope was equipped with an Apochromat 100×/1.49 numerical aperture oil-immersion objective lens and solid-state lasers with wavelengths of 488 nm, 561 nm, and 640 nm. The output powers at the fiber end were set to 15 mW for each laser. The images were captured using Nikon NIS-Elements software, specifically the 512 × 512 resolution setting. Z-stacks were acquired with a step size of 0.2 μm. The raw images were then reconstructed and processed using NIS-Elements AR Analysis software (version AR5.11.00 64bit).

The green channel (MTG, Rh123, GFP) was imaged using ’N-SIM: SIM488’ optical configuration, where its green emission was harvested through a high-rejection bandpass filter with a strict central wavelength of 526.5 nm. Meanwhile, the red channel (mCherry, MT-1) was scanned independently using the ’N-SIM: SIM561’ optical configuration, with its red emission captured through a corresponding bandpass filter centered at 605.5 nm. The third fluorophore channel (MTDR, CellROX DeepRed) was sequentially acquired using the ’N-SIM:SIM640’ optical configuration with a far-red emission window centered at 700.5 nm.

Cells were seeded onto glass-bottomed culture dishes (MatTek; P35G-1.5-14-C) and allowed to adhere for 24 hours. Subsequently, staining with commercial dyes was conducted for a duration of 30 min. Both MTG and MTDR were prepared as 1 mM stock solutions in DMSO and diluted in complete culture medium to a final concentration of 50 nM for live-cell staining. Before imaging, the cells were washed three times with PBS. During illumination, a self-made blue light LED array with an intensity of 300 μW/cm^2^ was employed. The subsequent analysis of the imaging data was carried out using ImageJ software.

### Mitochondria Contact Ratio Quantification by SIM

Super-resolution SIM images of living cells were acquired with mitochondria visualized using MTG, MTDR, or TOM20-EGFP, as indicated. Individual mitochondria were identified manually based on continuous mitochondrial profiles and morphology. The identification of two mitochondria as distinct objects was made independently of the distance criterion used to classify close apposition. For each pair of neighboring mitochondria, the shortest edge-to-edge distance between the two mitochondrial profiles was measured in reconstructed SIM images. A fluorescence-defined close apposition was classified when this distance was ≤ 4 pixels (≤ 124 nm under the present imaging conditions; 31 nm/pixel). A mitochondrion was classified as contact-positive if it participated in at least one such apposition. Each mitochondrion was counted once regardless of whether it contacted one or multiple neighboring mitochondria.

For each cell, contact ratio was calculated as:

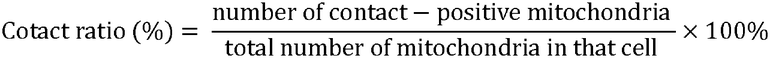

### TEM contact and nearest-neighbor analyses

TEM images were acquired using a Hitachi HT-7800 transmission electron microscope at 80 kV. Mitochondrial outer-membrane boundaries were identified manually. Two mitochondria were classified as forming a TEM-defined close contact when the shortest outer-membrane-to-outer-membrane distance was < 25 nm. Contact ratio was calculated separately for each cell as the number of mitochondria participating in at least one < 25 nm contact divided by the total number of mitochondria.

For the nearest-neighbor analysis, the shortest outer-membrane-to-outer-membrane distance between each mitochondrion and its nearest distinct neighboring mitochondrion was measured. The full distribution of individual mitochondrial nearest-neighbor distances is shown in Figure 1e. Statistical analysis was performed using nested t test.

### Toxicity Test

The CCK-8 assay was used to assess cell viability following blue light exposure. A cell suspension (100 μL, containing 1,000 cells/well) was dispensed into a 96-well plate. After a 24-hour pre-incubation, the entire plate was irradiated with blue light for a specified duration, with the control group covered with aluminum foil. Subsequently, 10 μL of CCK-8 solution was added to each well, and the plate was incubated for an additional 1 hour. Absorbance at 450 nm was measured to determine cell viability.

### Mitochondrial Morphology Quantification

Mitochondria were labeled with MTG or MTDR. After acquiring images using SIM, fluorescent images were imported into ImageJ. The image type was converted to 8-bit, and the threshold was auto adjusted. The area, shape descriptors, and threshold settings were refined using the “Set Measurements” function. Quantitative results were obtained through the “Analyze Particles” function. The “Aspect Ratio (AR)” value, calculated as the ratio of length to width (L/W), was recorded. Mitochondrial morphology was categorized into four groups based on AR values: Hyperfused (AR ≥ 5.0), Tubular (2.0 ≤ AR < 5.0), Intermediate (1.5 ≤ AR < 2.0), and Round (1.0 ≤ AR < 1.5).

### Oxygen Consumption (OCR) Rate Assay

To assess the mitochondrial OXPHOS function in the presence of blue light exposure, the OCR of cells expressing CRY2PHR-mCherry-Miro1TM was measured using a Seahorse XFe-96 Analyzer (Agilent Technologies, USA). The OCR measurements were conducted in real time. Cells were divided into two groups: one group for blue light illumination and another group kept in the dark. Additionally, cells expressing CRY2PHR-mCherry-Miro1TM were also divided into two groups: one group for blue light illumination and another group kept in the dark. These cells were seeded into XFe-96 cell culture plates (Agilent Technologies, USA) at a density of 0.8 × 104 cells per well, using DMEM supplemented with 10% FBS. After a 48-hour incubation period, the culture medium was aspirated and replaced with pre-warmed unbuffered “Seahorse medium” (XF DMEM) at a pH of 7.4. The Seahorse medium was supplemented with 1 mM sodium pyruvate, 10 mM glucose, and 2 mM L-glutamine. Prior to the measurement, one group of cells and one group of cells expressing the plasmid were subjected to blue light exposure at an intensity of 300 μW/cm^2^. Subsequently, the OCRs of the cells were evaluated using the XF Cell Mito Stress Test Kit (Agilent Technologies, USA), following the instructions provided by the manufacturer. In certain cases, the cells were subjected to treatment with specific compounds from Agilent Technologies, including 1 μM oligomycin A (an electron transport inhibitor), 2 μM FCCP (an electron chain transport accelerator), and 500 nM rotenone (a complex I inhibitor) with 1 μM antimycin A (a complex III inhibitor).

### ATP Quantification Assay

ATP quantification was performed using the Invitrogen™ ATP Determination Kit. Two experimental groups, each consisting of three independent samples, were prepared: control (not transfected) group and transfected (CRY2PHR-mCherry-Miro1TM) group. Cells were seeded overnight into Costar 96-well plates (Corning, USA) for adherence. Transfection was then performed for transfected group, followed by a 24-hour incubation for plasmid expression. After a certain time of blue light exposure, the samples were harvested and processed according to the manufacturer’s protocol of the ATP Determination Kit. Luminescence was measured using a microplate reader (BioTek Instruments, Inc., USA).

### Mitochondria Membrane Potential

Rhodamine 123, a cell-permeant, cationic, green-fluorescent dye, was used to assess local contact-associated MMP. in living cells, with fluorescence changes dependent on the membrane potential. Rhodamine 123 was added to the cell samples to achieve a final concentration of 1 μM, and staining was carried out for 30 minutes.

The second independently live-cell MMP at the complementary cell-level using MT-1 MitoMP Detection. Cells were washed and incubated with a 1:1000 dilution of MT-1 working solution at 37°C for 30 min in a 5% CO^2^ incubator.

Cells were washed three times with PBS and kept in complete medium before imaging. MT-1 and mCherry were detected in the same red acquisition channel. For each treatment condition, we calculated the ratio of fluorescence intensity between transfected and control cells and compared these ratios across conditions. Because the measured fluorescence could include contributions from mCherry, these ratios were interpreted as relative changes in red-channel fluorescence rather than direct, uncontaminated measurements of MMP.

### Mito-contact Sites Identification and MMP Calculation

A custom image analysis algorithm was developed for study light induced mito-contact sites. The process begins by automatically detecting protein aggregates as distinct red puncta in the red fluorescence channel, accomplished by applying a high-percentile intensity threshold to identify the brightest local maxima. Following detection, a circular region of interest with a 10-pixel radius is defined around the center of each punctum, designated as the “mito-contact site.” To ensure the analysis is restricted to mitochondria, a binary mask is generated from the corresponding green fluorescence channel. The algorithm then calculates the mean green fluorescence intensity (representing MMP) within the mitochondrial pixels of each mito-contact site and compares this value to the average intensity of the background, which is defined as all mitochondrial areas located outside of these specific sites. The code could be found in https://github.com/eugenelet/Modulating-inter-mitochondrial-contacts-to-increase-membrane-potential.

### ROS measurement

Intracellular ROS was assessed using CellROX Deep Red under acute 20 min blue light activation and prolonged 3 h blue light exposure. Cells were incubated with CellROX Deep Red at 2.5 μM for 30 min in 37°C, 5% CO^2^ incubator. Integrated CellROX fluorescence was quantified within manually defined cell ROIs.

### RNA-Sequencing and Data Analysis

Two groups (control and transfected), with each of six dishes, were prepared. Three for dark and three for blue light treatment ensured three independent repeats for each condition. HeLa cells were seeded to 60 mm dish and incubated for 24 h for adhering. Did transfection process for transfected group samples and waiting for 24 h for good expression. The blue light treatment group were exposure under blue light for a certain time before it was fixed. Then, all dishes were washed with cold PBS three times and fixed with lysis buffer for RNA extraction. Total RNA extraction and directional polyA RNA-seq were performed by the Genomics, Epigenomics and Sequencing Core at the University of Cincinnati.

The RNA quality was assessed using a Bioanalyzer (Agilent, USA), confirming high RNA quality. PolyA RNA was enriched using the NEBNext Poly(A) mRNA Magnetic Isolation Module (New England BioLabs, USA) combined with the SMARTer Apollo automated NGS library prep system (Takara Bio USA, USA) using 1 μg of total RNA as input. Library preparation was carried out using the NEBNext Ultra II Directional RNA Library Prep Kit (New England BioLabs, USA) with 8 PCR cycles. Following library quality control via Bioanalyzer and quantification using real-time qPCR (NEBNext Library Quant Kit, New England BioLabs, USA), individually indexed libraries were pooled proportionally and sequenced using the NextSeq 550 sequencer (Illumina, USA). Approximately 25 million reads per sample were generated. Fastq files for downstream data analysis were automatically generated via the Illumina BaseSpace Sequence Hub following sequencing. Differential expression analysis was conducted using the DESeq2 R package. Genes were considered significantly differentially expressed if the adjusted p-values (q-values) were less than 0.05. Mitochondrial genes were screened out by using Human MitoCarta3.0: 1136 mitochondrial genes. Gene set enrichment analysis (GSEA) was performed using the GSEApy Python package. We used pre-ranked gene expression data to assess overrepresented Gene Ontology (GO) categories and Kyoto Encyclopedia of Genes and Genomes (KEGG) categories. Specially, we analyzed three GO categories – Biological Process, Molecular Function, and Cellular Component – from GO gene subsets respectively; and also analyzed KEGG pathway for the wider range of biochemical processes.

*C. elegans* Experiments

Strains and Maintenance: The animal experiments were approved by the Cincinnati Children’s Hospital Medical Center Institutional Biosafety Committee. Nematode growth medium (NGM) was used for *C. elegans* culture and all maintenance and experiments were carried out at 20 °C. OP50 Escherichia coli was used as a food source for all experiments. Egg-lay-synchronized hermaphrodite animals were used for all experiments at the stage noted. Strain ZAC307 cchEx307[pTG028_eft-3p::CRY2::-mCherry-Miro1TM + myo-2p::RFP] expresses CRY2-mCherry-Miro1TM in all cells. ZAC307 was created by injecting N2 Young adult worms with a mix containing 0.25 mM EDTA, 2.5 mM Tris-Cl, 10 ng µl−1 pTG028[eft-3p::CRY2::-mCherry-Miro1TM] plasmid, 2.5 ng µl−1 myo-2p::RFP plasmid, 90 ng µl−1 NEB 1kB ladder. Offspring were selected for the presence of the transgenic array based the red pharyngeal myo-2p::RFP marker. The expression of CRY2::-mCherry-Miro1TM was verified in L4 worms with a Nikon A1R Confocal at 60× using the 60× Apo λ S Oil Immersion Objective.

Lifespan Analysis: For the lifespan study, four groups of *C. elegans* were used. On day 0, L4 animals were transferred to 3 × 60 mm NGM plates, 20 animals per plate (n = 60). For the Control group, N2 (wild-type) animals were kept in a box, in darkness at 20 °C. For the Control + light group, N2 (wild-type) animals were kept in a blue light box (300 μW/cm^2^) at 20 °C. For the Transgenic group, array positive ZAC307 animals were kept in a box, in darkness at 20 °C. For the Transgenic + light group (optogenetic treatment), array positive ZAC307 animals were kept in the blue light box (300 μW/cm^2^) at 20 °C. Every two days, each set of 20 adult animals was moved to a new 60-mm NGM plate to leave any progeny behind, and the number of living animals was assessed. Animals were counted as living if they moved or responded to stimuli when they were picked or tapped.

### Data Analysis

Statistical analyses were performed using Prism 8 (GraphPad). Two-group comparisons were performed using two-tailed Student’s t tests where indicated. MT-1 transfected-to-control fluorescence ratios across treatment conditions were analyzed using one-way ANOVA followed by Tukey’s post hoc test. Survival curves were compared using the log-rank (Mantel-Cox) test.

For imaging analyses involving multiple mitochondrial measurements within the same cell, individual mitochondria were not treated as independent biological replicates. Instead, to avoid pseudo replication, measurements from multiple mitochondria within a single cell were averaged to represent a single cell-level or single image data point (n). The total number of cells/images pooled from 3 independent experimental trials, along with the specific statistical tests, are detailed in the corresponding figure legends. Statistical analysis and graphing were performed using Prism 8 software (GraphPad) or Excel (Microsoft). Image analysis was carried out using ImageJ-win64 software (NIH Image). The final assembly of images was done using PowerPoint (Microsoft).

### Statistics and Reproducibility

Statistics were performed in Prism 8 (GraphPad). No data was excluded from the analysis. No statistical method was used to pre-determine sample size. Within experimental groups, samples were randomized for each experimental replicate. The investigators were not blinded to allocation during experiments and outcome assessment. Data distribution was assumed to be normal, but this was not formally tested, therefore data distributions are visualized in each figure.

## Data Availability

All data supporting the findings of this study are available either in the article and/or its Supplementary Information files. Source data are provided with this paper.

