## Supplemental Figures for "Modulating inter-mitochondrial contacts to increase membrane potential for mitigating blue light damage"

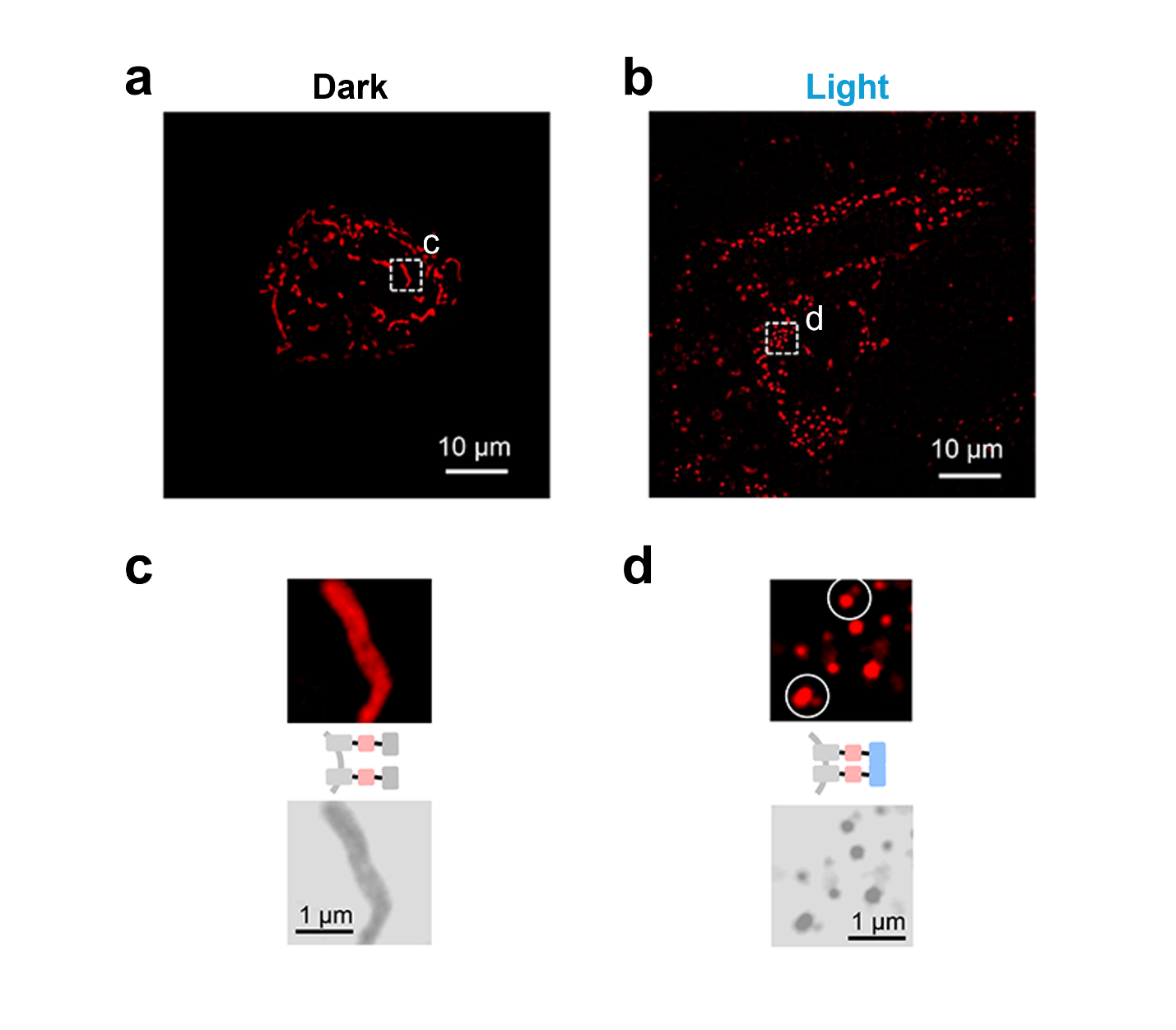

**Figure S1.** Representative SIM images of HeLa cells expressing CRY2PHR-mCherry-Miro1TM.

**(a)** 0 min (dark) and **(b)** 10 min of blue light exposure at 300 μW/cm^2^.

**(c,d)** Schematic representation of the expressing proteins on mitochondria and the zoom-in view of marked region of interests in (a) and (b), respectively. For better comparison between the changes of fluorescence, the corresponding gray images for the enlarged images were shown. The circles in (d) indicate the protein puncta.

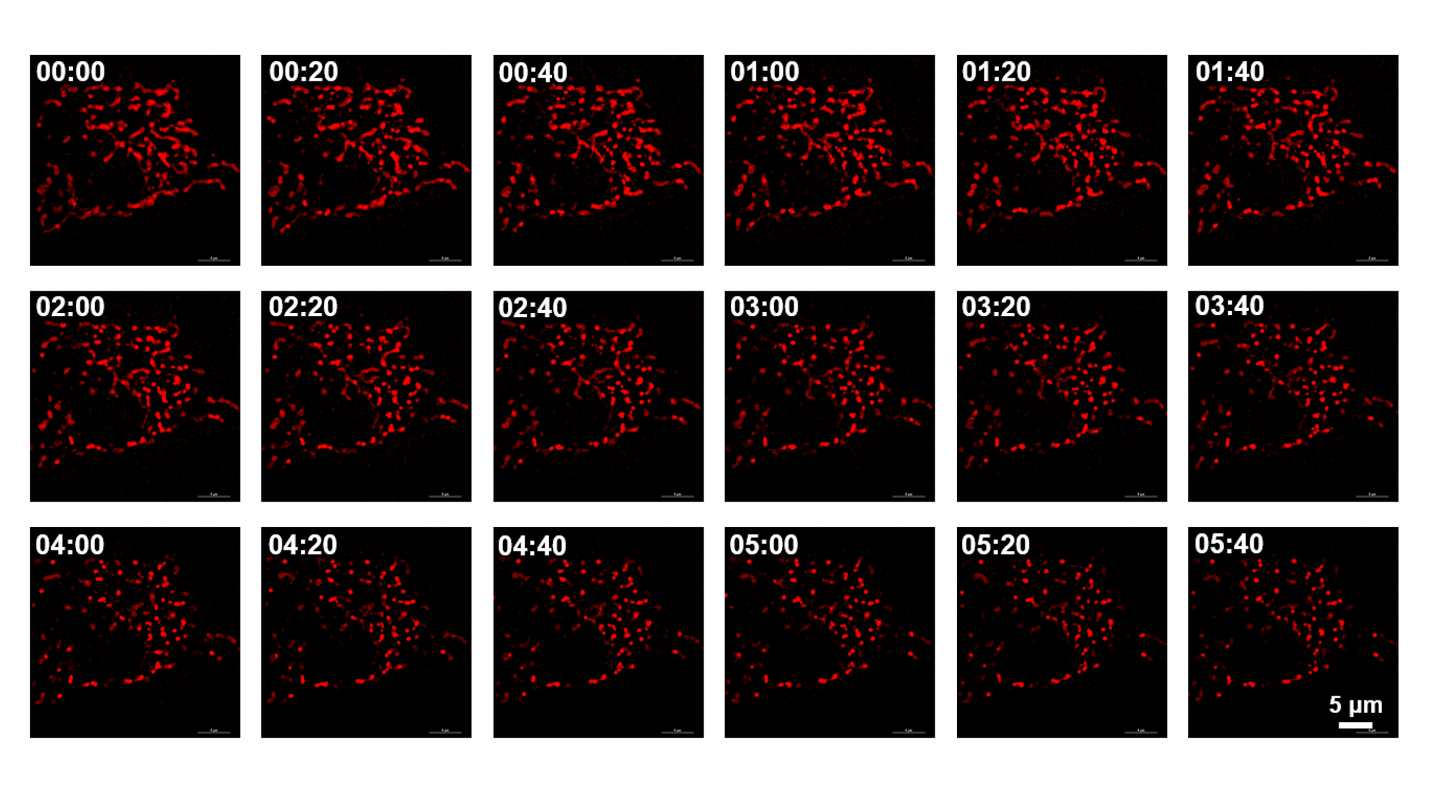

**Figure S2.** Time-stamped SIM images of optogenetically induced clustering of CRY2PHR in live HeLa cells. All images shared the same scale bar. Time shown in minutes and seconds.

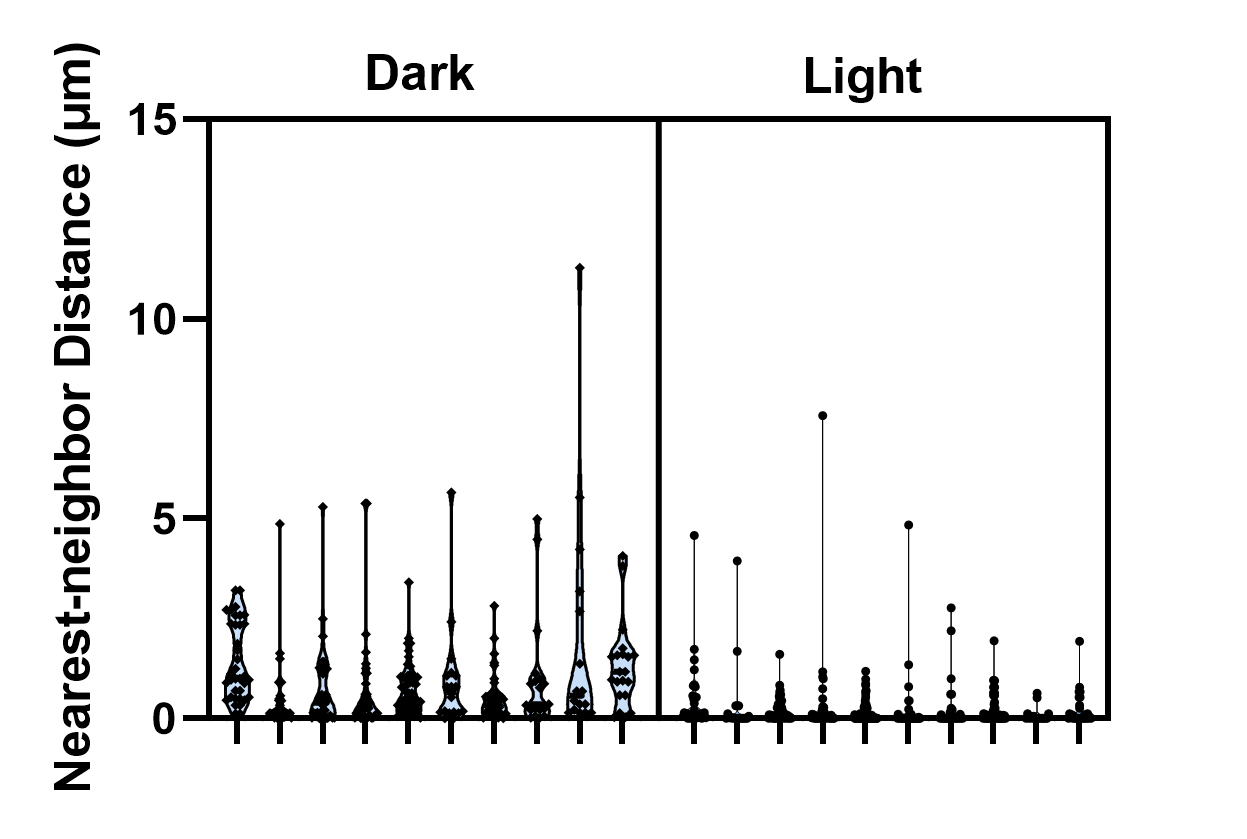

**Figure S3.** Violin plots showing nearest-neighbor distance of mitochondria in Dark vs. Light groups. Each violin plot represents the spatial data quantified from a single cell in TEM images.

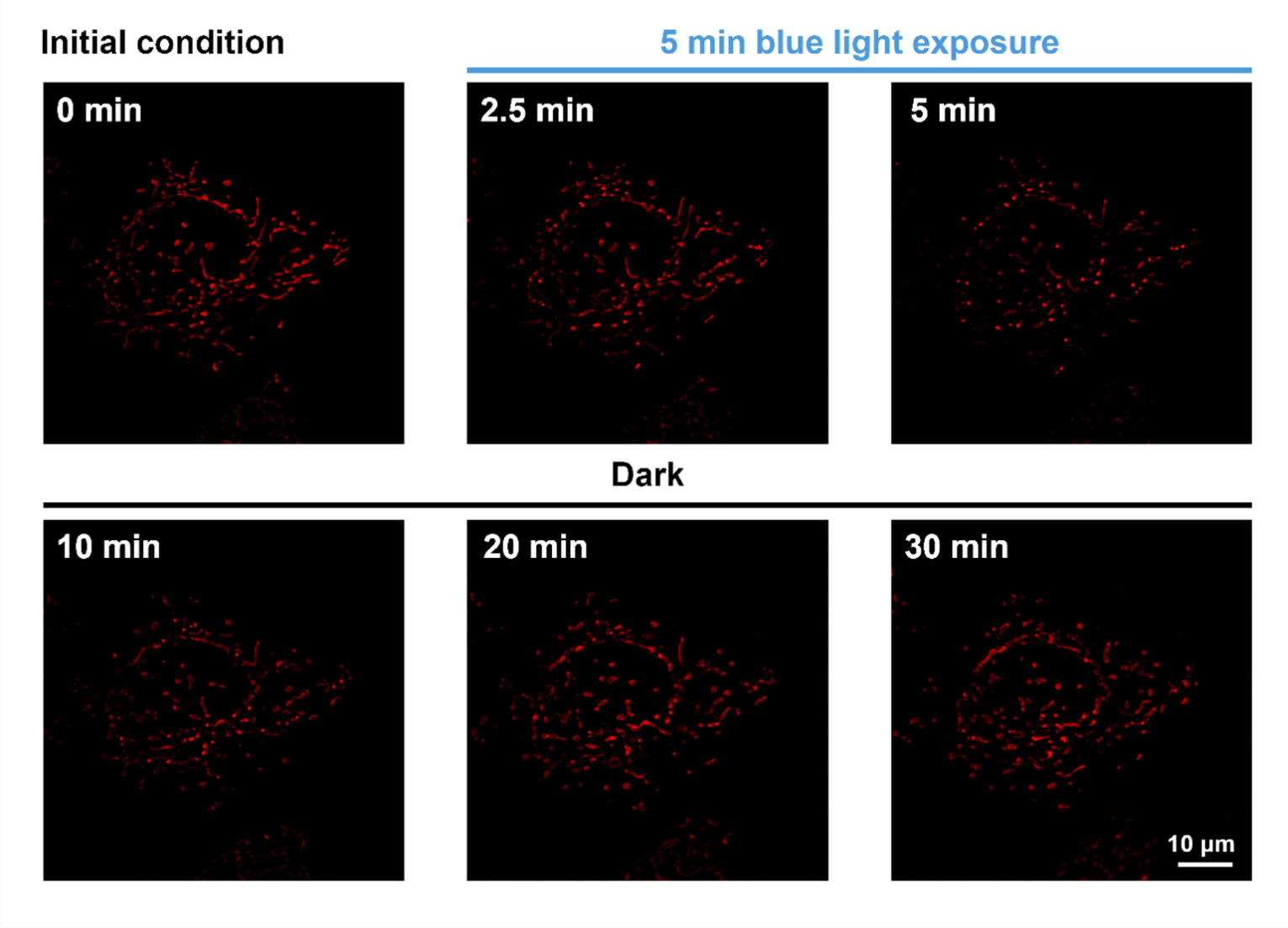

**Figure S4.** The time-lapse SIM images of light-induced CRY2 oligomerization and dark dissociation.

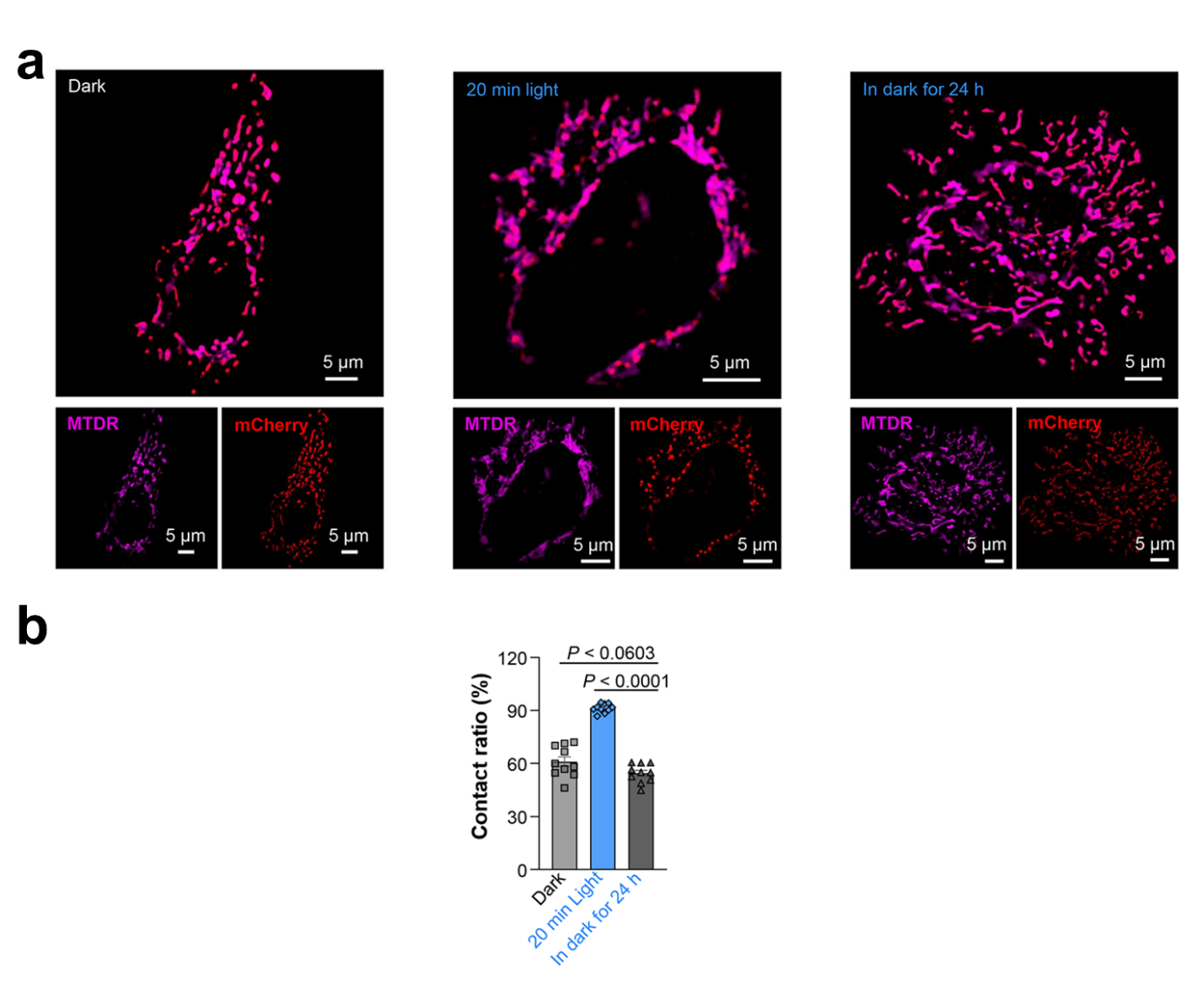

**Figure S5.** The reversibility of mitochondrial clustering by the optogenetic system.

**(a)** The SIM images of living HeLa cells expressing CRY2PHR-mCherry-Miro1TM and stained with MTDR kept in the dark (left), immediately after 20 min blue-light exposure (300 μW/cm^2^) (middle), and illuminated cells following 24 h dark incubation (right).

**(b)** The percentage of contact ratio in HeLa cells expressing CRY2PHR-mCherry-Miro1TM and stained with MTG/MTDR in each condition. Data are given as *M* ± *SEM* (*n* = 10).

Statistical differences between the experimental groups were analyzed using a double-tailed Student’s *t* test. All *P* values less than 0.05 were considered to indicate statistical significance.

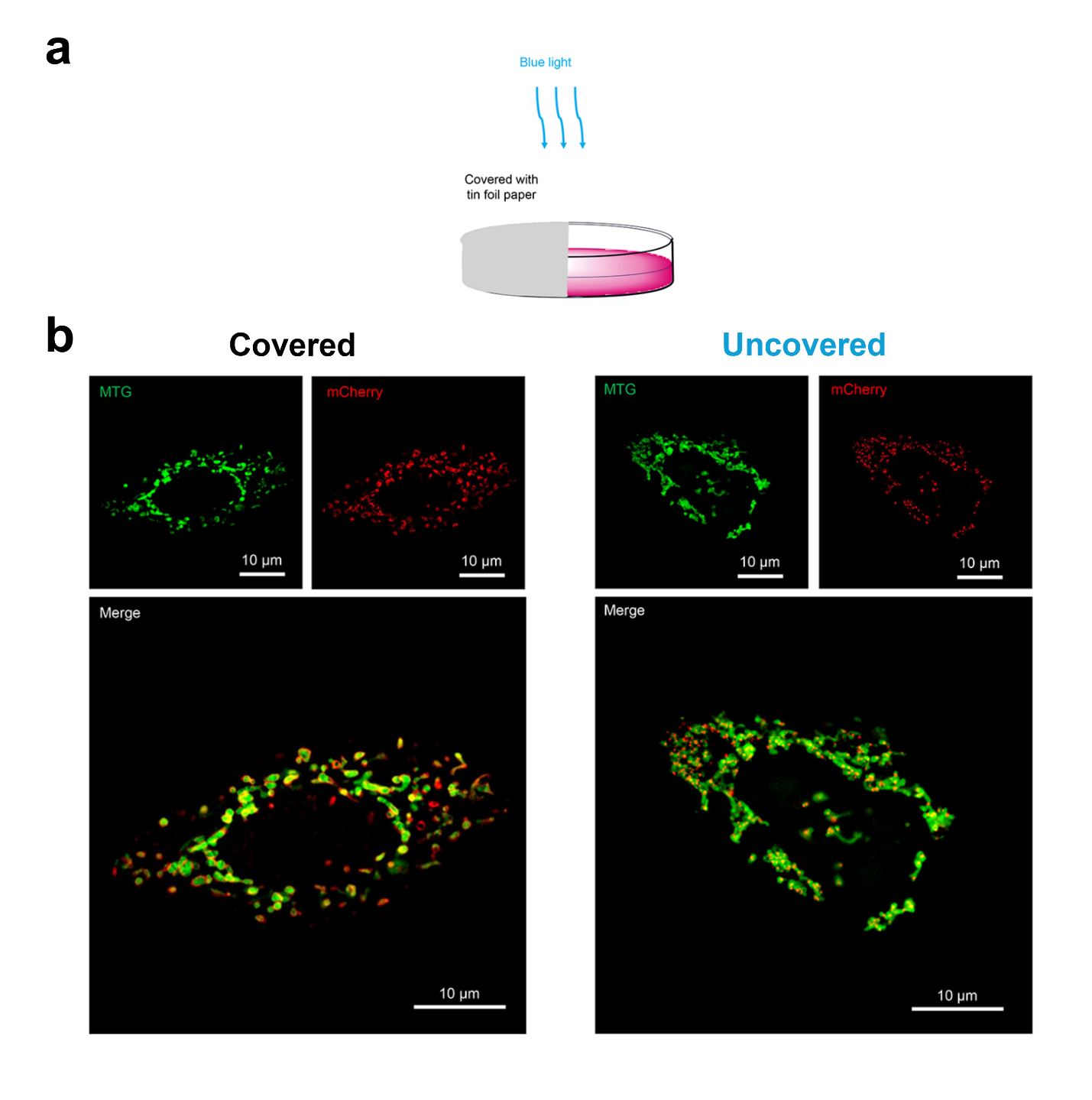

**Figure S6.** Spatial control of the optogenetic system.

**(a)** Schematic representation of the spatial control experiment, where half of the culture dish was covered with tin foil paper to darken that portion of the dish.

**(b)** The SIM images of HeLa cells expressing CRY2PHR-mCherry-Miro1TM and stained with MTG by spatial control with or without light exposure for 20 min.

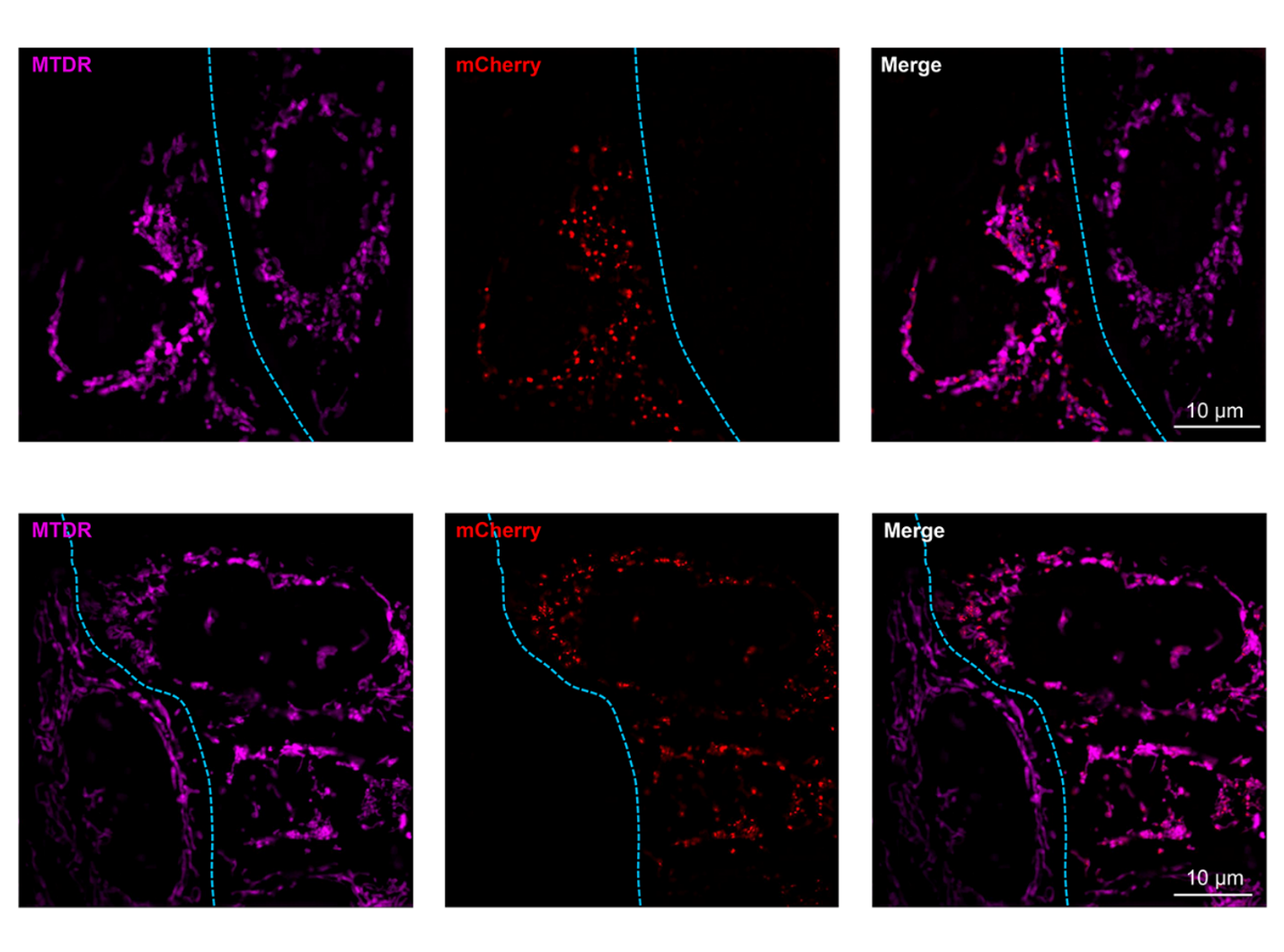

**Figure S7.** The photoactivatable protein is important for the optogenetic system.

The SIM images of living HeLa cells expressing CRY2PHR-mCherry-Miro1TM and adjacent untransfected cells and stained with MTDR under blue light for 20 min. All cells were cultured in the same dish.

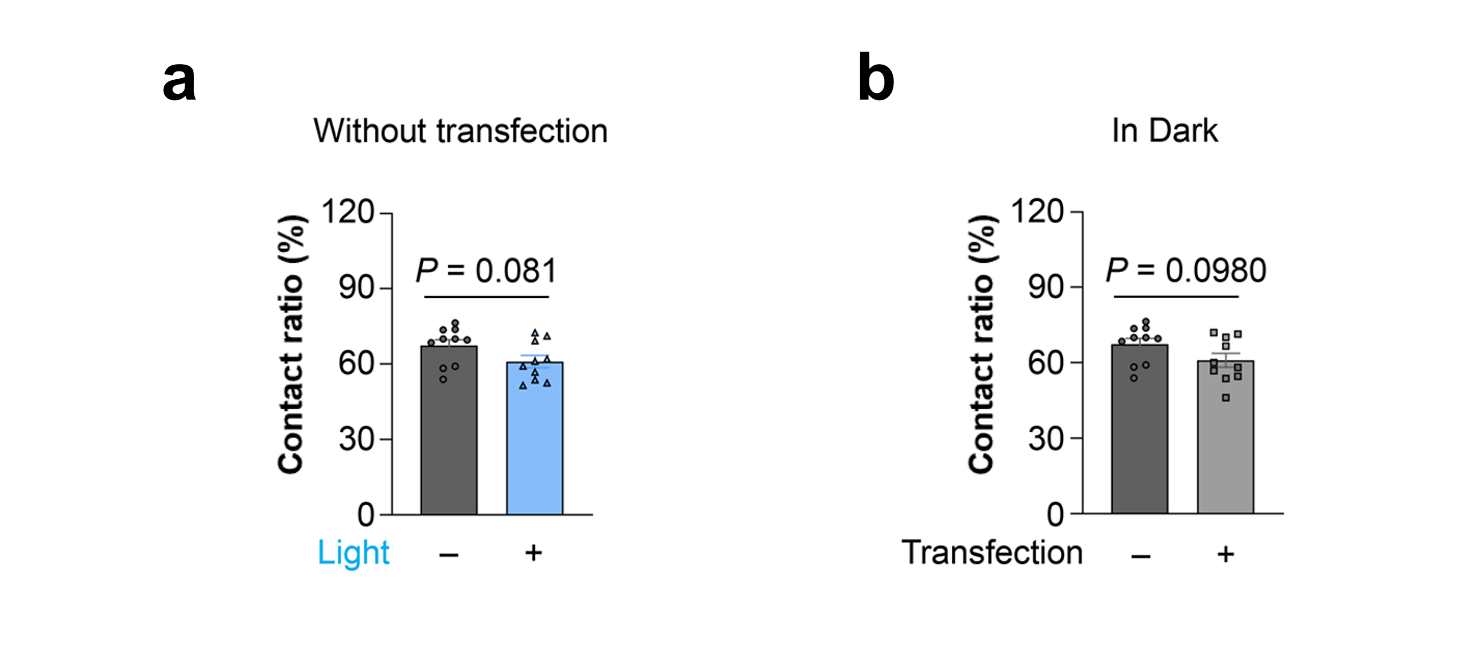

**Figure S8.** The photoactivatable protein and light are the essential conditions for the optogenetic system.

**(a)** Quantitative analysis of the percentage of mitochondria having one or more inter-contact sites to other mitochondria for HeLa cells stained with MTDR without (Dark) or with (Light) blue light exposure for 20 min.

**(b)** The comparison between the cells in the dark without and with transfection of CRY2PHR-mCherry-Miro1TM.

Data are given as *M* ± *SEM* (*n* = 10). Statistical differences between the experimental groups were analyzed using a double-tailed Student’s *t* test. All *P* values less than 0.05 were considered to indicate statistical significance.

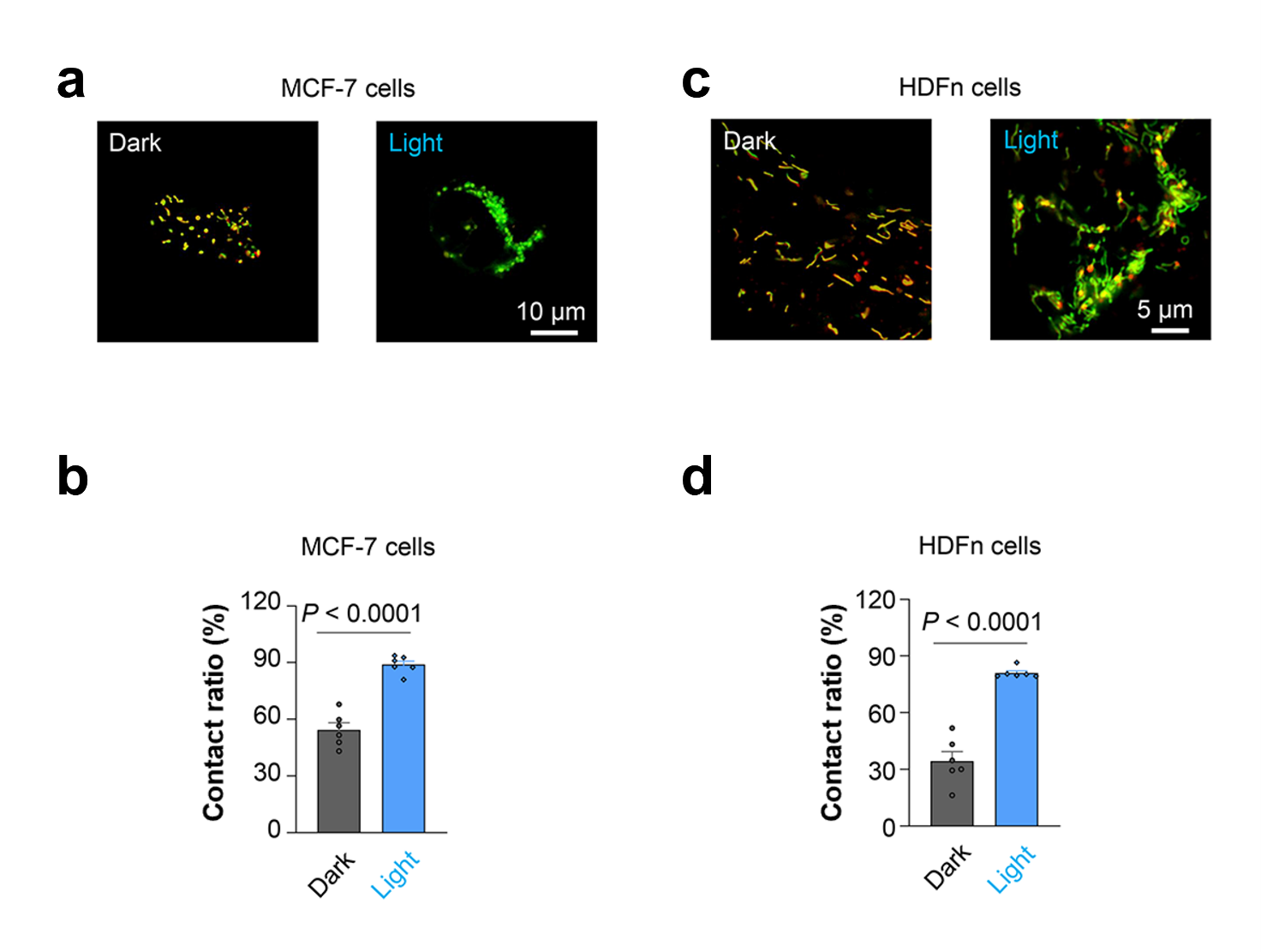

**Figure S9.** The optogenetic mito-contact system in MCF-7 and HDFn cells.

**(a)** SIM images of living MCF-7 cells expressing CRY2PHR-mCherry-Miro1TM and stained with MTG in dark or under blue light for 20 min.

**(b)** Quantitative analysis of contact ratio for MCF-7 cells expressing CRY2PHR-mCherry-Miro1TM and staining MTG in dark or with blue light exposure.

**(c)** SIM images of living HDFn cells expressing CRY2PHR-mCherry-Miro1TM and stained with MTG in dark or under blue light for 20 min.

**(d)** Quantitative analysis of contact ratio for HDFn cells expressing CRY2PHR-mCherry-Miro1TM and staining MTG in dark or with blue light exposure.

Data in (b) and (d) are given as *M* ± *SEM* (*n* = 6). Statistical differences between the experimental groups were analyzed using a double-tailed Student’s *t* test. All *P* values less than 0.05 were considered to indicate statistical significance.

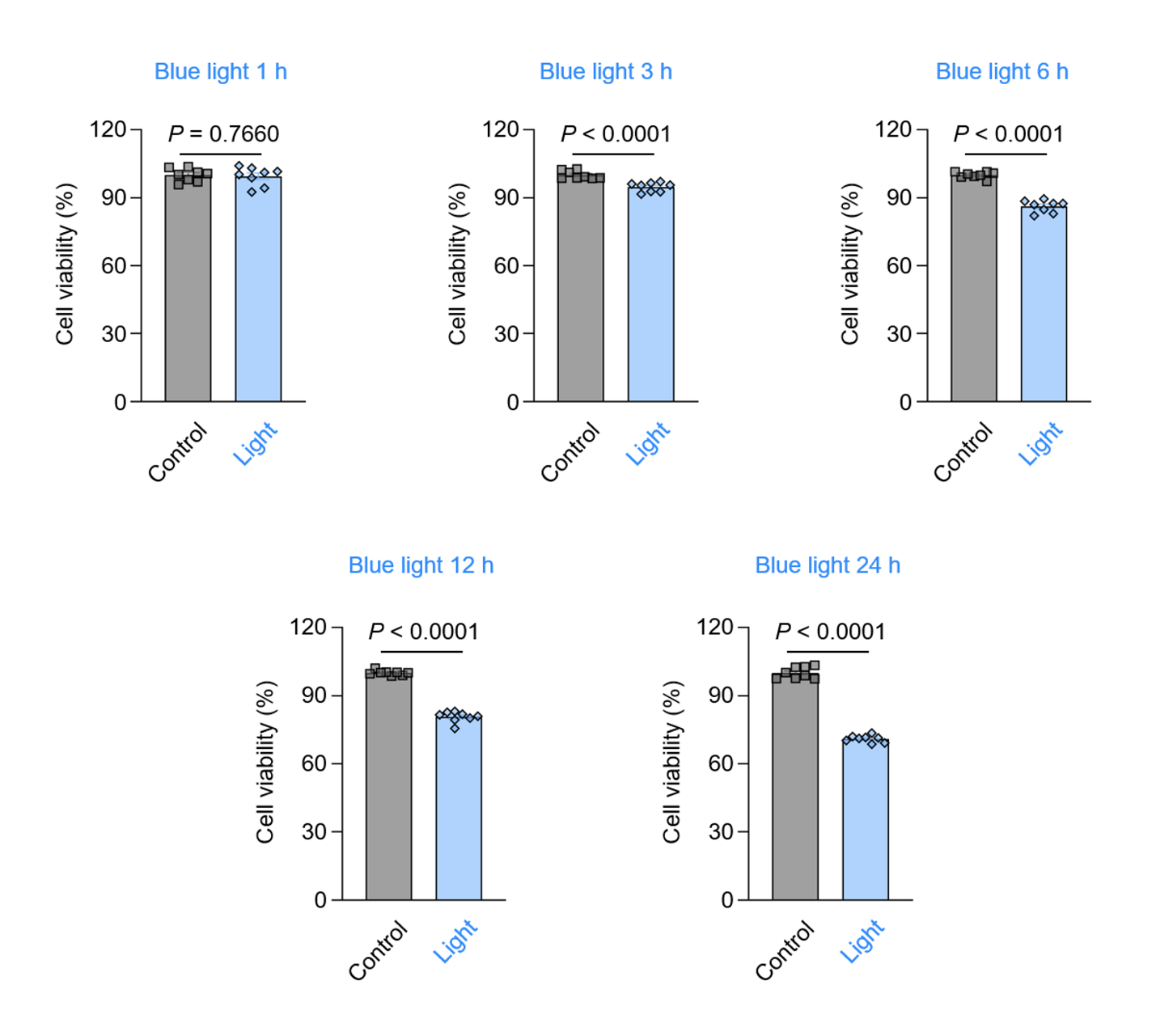

**Figure S10.** The phototoxicity of blue light to HeLa cells.

The viability of HeLa cells with blue-light exposure for different times. Data are given as *M* ± *SEM* (*n* = 8). Statistical differences between the experimental groups were analyzed using a double-tailed Student’s *t* test. All *P* values less than 0.05 were considered to indicate statistical significance.

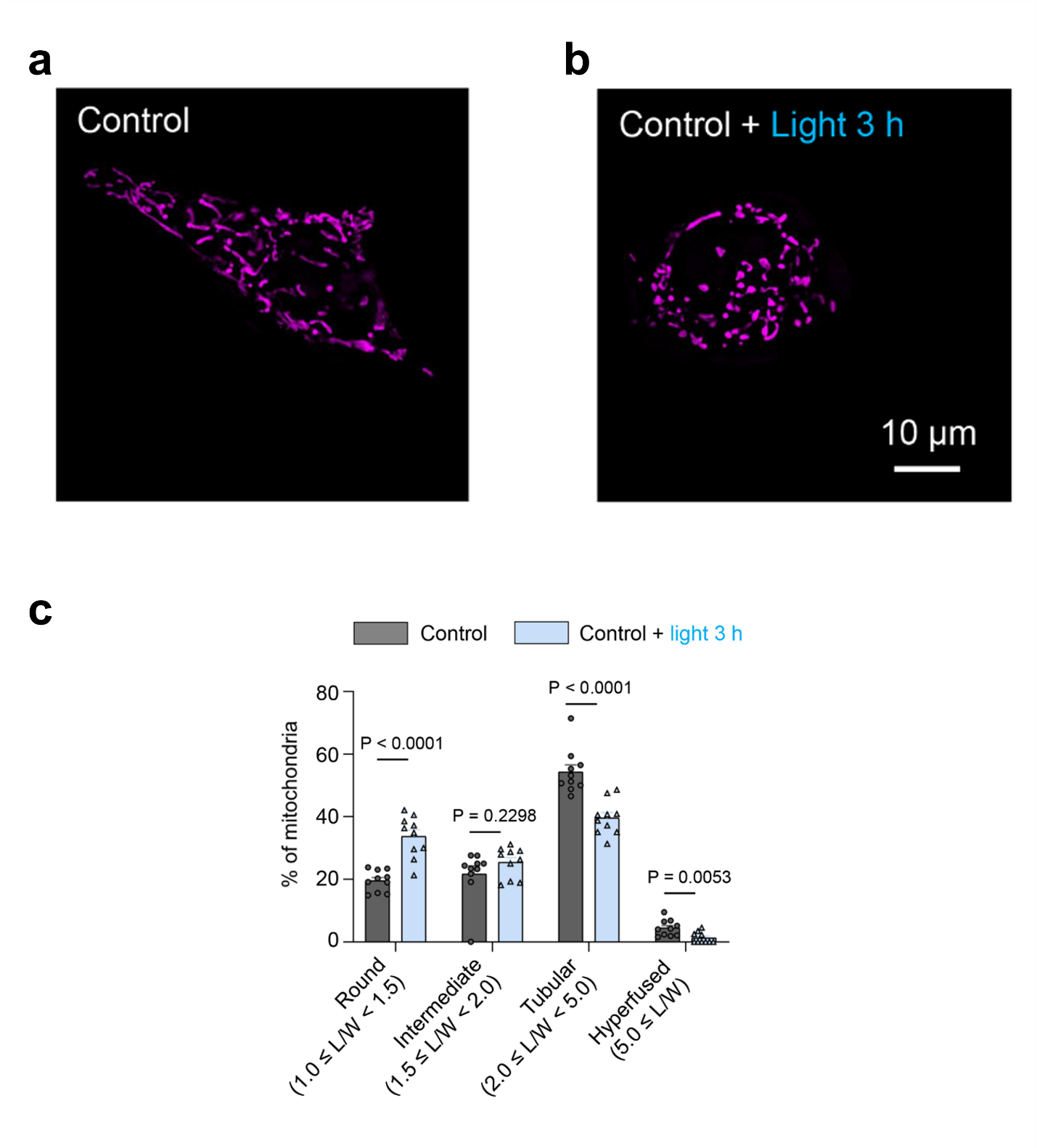

**Figure S11.** Mitochondria are damaged by blue light after 3 h exposure.

**(a,b)** The SIM images of MTDR stained HeLa cells with or without blue light exposure for 3 h.

**(c)** Quantitative analysis of mitochondrial morphology in (a,b).

Data in (c) are given as *M* ± *SEM* (*n* = 10). Statistical differences between the experimental groups were analyzed using a double-tailed Student’s *t* test. All *P* values less than 0.05 were considered to indicate statistical significance.

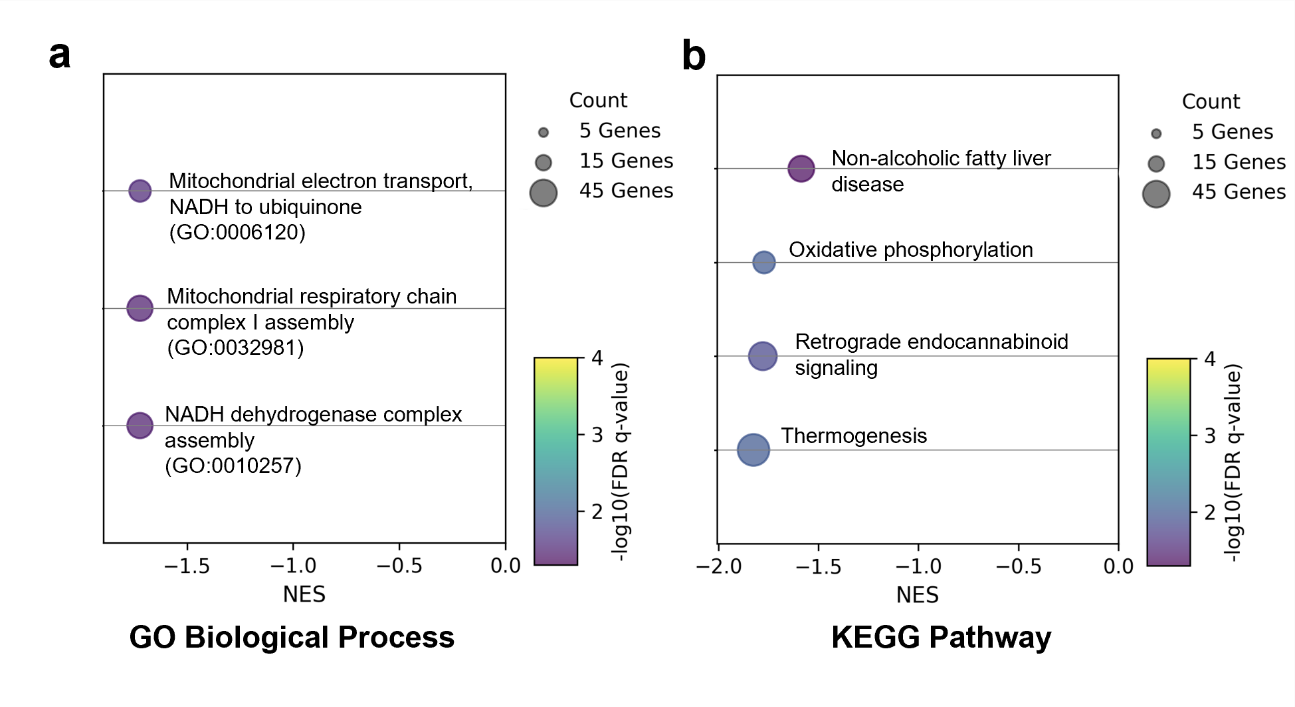

**Figure S12.** Mitochondrial related genes enrichment analysis.

**(a)** Bubble plot for Gene Ontology (GO) biological process enrichment analysis of mitochondrial related differential genes from Control group.

**(b)** Kyoto Encyclopedia of Genes and Genomes (KEGG) pathway enrichment analysis of mitochondrial related differential genes from Control group.

In the bubble plots, the x-axis shows Normalized Enrichment Score (NES), and y-axis shows pathway terms. The size of each circle represents the number of genes. The color of the circles presents various q-values. All terms presented above have q values less than 0.05 that indicate significance. The mitochondrial related differential genes from Transfected group have no significant terms in GO or KEGG pathway analysis.

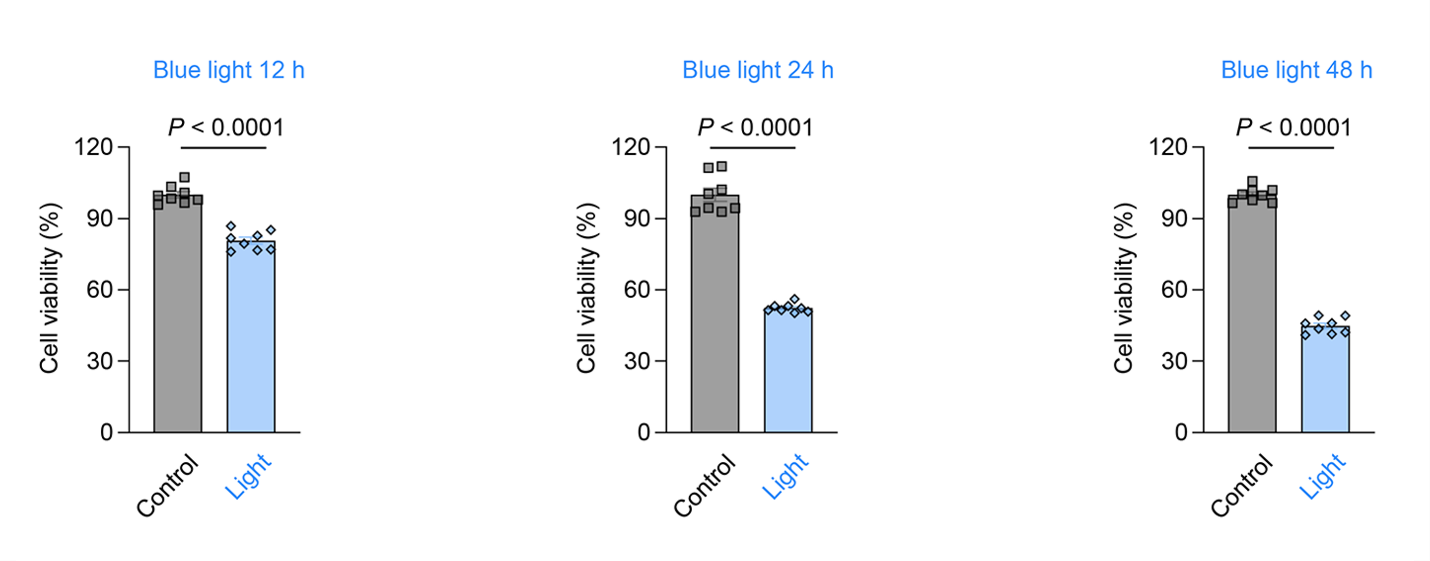

**Figure S13.** The phototoxicity of blue light to ARPE-19 cells.

The viability of ARPE-19 cells with blue-light exposure for different times. Data are given as *M* ± *SEM* (*n* = 8). Statistical differences between the experimental groups were analyzed using a double-tailed Student’s *t* test. All *P* values less than 0.05 were considered to indicate statistical significance.
